# Smooth muscle LRRC8A knockout reduces O_2_^·-^ influx, inflammation, senescence and atherosclerosis

**DOI:** 10.64898/2025.12.01.691680

**Authors:** Sourav Panja, Hyehun Choi, Hong N. Nguyen, Julia K. Russolillo, Sergey Dikalov, Ryan J. Stark, Fred S. Lamb

**Affiliations:** Department of Pediatrics, Division of Critical Care, Vanderbilt University Medical Center, Nashville, TN 37232; Department of Internal Medicine, Division of Clinical Pharmacology, Vanderbilt University Medical Center, Nashville, TN 37232

**Author notes:** Corresponding author: Fred S. Lamb MD, PhD, Vanderbilt University Medical Center Department of Pediatrics, 2215 Garland Avenue, Light Hall-1055D Nashville, TN 37232-3122. These authors contributed equally to this work (Sourav Panja, Hyehun Choi).

**Keywords:** Leucine Rich Repeat Containing 8, vascular smooth muscle, inflammation, oxidative stress, senescence, atherosclerosis

## Abstract

**Background:** Leucine Rich Repeat Containing 8A (LRRC8A) anion channels (VRACs) associate with NADPH oxidase 1 (Nox1) and support extracellular superoxide (O_2_**^·^**^-^) production, inflammation, and contractility in vascular smooth muscle cells (VSMCs). We proposed previously that VRACs also support influx of O_2_**^·^**^-^ to localize cytoplasmic redox signals.

**Methods:** We assayed O_2_**^·^**^-^ influx, assessed changes in mRNA expression (RNAseq), and tested multiple phenotypes of cultured LRRC8A knockout (KO) VSMCs. Aortic atherosclerotic burden and inflammation, and mesenteric vascular reactivity were compared between wild type (WT) apolipoprotein E null (ApoE^-/-^) mice and VSMC-specific LRRC8A KO, ApoE^-/-^ mice.

**Results:** KO cells were less permeable to extracellular O_2_**^·^**^-^, produced less mitochondrial O_2_**^·^**^-^, and experienced less oxidant stress (GSH/GSSG, lipid peroxidation, Nrf2 activity) than WT. RNAseq and reporter assays demonstrated reduced pro-inflammatory (NF-κB and Hif1α) transcription. KO cells also under-expressed multiple senescence markers and had longer telomeres. Inflammation causes a metabolic shift from oxidative phosphorylation (OCR) to glycolysis (ECAR). Both pathways were less active in KO cells, as was expression of glycolytic enzymes. Mitochondrial membrane potential was lower in KO cells, but ADP/ATP and NADP^+^/NADPH were unaltered, suggesting lower energy demand. Consistent with this, proliferation and migration were both reduced in KO cells. Protein-protein interaction analysis of RNAseq data (PPI Hub) identified Epidermal Growth Factor Receptor (EGFR) signaling. Resting phosphorylation of EGFR (pY1068) and AKT (pS473 and pT308) were all reduced in KO cells, as was and EGF-induced pY1068 and pS308. Following 15 weeks of exposure to a high fat diet (42%) VSMC-specific LRRC8A^-/-^ and 8A^+/-^, ApoE^-/-^ mice had reduced atherosclerotic lesion area, aortic senescence (β-Gal), inflammation (ICAM, VCAM) and proliferation marker (PCNA) expression compared to WT, ApoE^-/-^ controls. Mesenteric artery vasomotor function was also preserved, and the abundance of MYPT1 and CPI17 was lower in KO vessels. Uptake of oxidized LDL (OxLDL) was significantly reduced in KO VSMCs and impaired by VRAC block in WT cells.

**Conclusions:** Loss of LRRC8A reduced O_2_**^·^**^-^ influx, oxidative stress, inflammation and senescence, lowers energy demand, and in the setting of hypercholesterolemia impaired uptake of oxidized LDL and abrogated atherosclerosis. VSMC VRACs may be novel targets of vascular anti-inflammatory therapy.

**Novelty and Significance:** *What Is Known?:*

- Chronically inflamed vascular smooth muscle cells (VSMCs) are critical drivers of atherosclerotic plaque development.
- In VSMCs, Nox1-derived superoxide drives redox signaling that promotes inflammation, phenotypic switching, and senescence.
- LRRC8A family volume-regulated anion channels (VRACs) physically associate with Nox1 and support TNFα-induced ROS production, receptor endocytosis, NF-κB activation, and proliferation.
- Hypercholesterolemia causes VSMC inflammation via oxidized LDL (OxLDL) activation of scavenger receptors (e.g., LOX-1, CD36) that stimulate oxidant production and EGFR/PI3K–Akt–NF-κB signaling.

*What New Information Does This Article Contribute?:*

- LRRC8A channels support influx of extracellular superoxide which impacts cytosolic redox status, metabolism and Nrf2, HIF-1α, and NF-κB activity.
- Loss of LRRC8A reprograms VSMCs into a low-oxidative-stress, low-energy-demand state, characterized by diminished proliferation and migration, attenuated senescence, and preservation of telomere length.
- Loss of LRRC8A inhibits EGF-stimulated EGFR and Akt phosphorylation and uncouples oxidant-dependent growth factor signaling from downstream inflammatory and metabolic remodeling.
- VSMC-specific LRRC8A knockout or heterozygosity in hypercholesterolemic ApoE⁻/⁻ mice reduces aortic inflammation and senescence, reduces atherosclerotic lesion burden, and preserves mesenteric artery vasomotor function.
- LRRC8A/VRAC activity is necessary for uptake of oxidized but not native LDL in VSMCs.
- LRRC8A is a novel regulator of scavenger receptor-dependent lipoprotein handling and a potential therapeutic target to uncouple oxidant signaling from VSMC lipid overload and inflammation in the vessel wall.

## INTRODUCTION

LRRC8 family genes (LRRC8A-E) encode volume-regulated anion channels (VRACs), originally identified based on activation by hypotonicity and their contribution to regulatory volume decreases. LRRC8A is obligatory for the trafficking of all VRACs to the plasma membrane^1^, where they have also evolved to serve other important roles^2–5^. In vascular smooth muscle cells (VSMCs) these channels are regulated by NADPH oxidase (Nox)-derived oxidants produced in response to cell swelling, and to signaling molecules including angiotensin II, TNFα and growth factors^6,7^. VRACs support proliferation in multiple cell types including VSMCs^8,9^, and the magnitude of their current varies with the cell cycle^10,11^, being more active in proliferative cells^12^. VRACs are required for TNFα-induced NF-κB activation^13,14^ and proliferation of VSMCs^15^. These signals require production of extracellular superoxide (O_2_**^·^**^-^) by Nox1 which physically-associates with LRRC8A, 8C and 8D, and oxidase activity depends upon VRAC function^16,17^.

VRACs are linked to redox balance in at least two ways^18^, they regulate oxidant-dependent signaling^16,17,19^, and their activity is modulated by oxidants^16,20^. We previously proposed that VRACs also provide a mechanism by which extracellular O_2_**^·^**^-^ created by Nox1 can enter the cytoplasm to drive localized redox reactions (See^19^ for review). This concept was supported by several indirect observations. TNFα receptor endocytosis and c-Jun N-terminal kinase activation were impaired by extracellular superoxide dismutase, while catalase did not affect these signals^17^. Furthermore, activation of apoptosis signaling-related kinase 1 (ASK1) by TNFα, which depends upon oxidation of cytoplasmic thioredoxin, was Nox1-dependent and potentiated by knockdown of extracellular SOD^19^. In addition, TNFα-induced RhoA activation and sulfenylation (oxidation) of the cytoplasmic LRRC8A-associated scaffolding protein Myosin Phosphatase Rho Interacting Protein (MPRIP), were both LRRC8A-dependent^21^. Recently this hypothesis was supported by direct evidence for LRRC8A-dependent O_2_**^·^**^-^ influx in neurons, where disruption of VRAC function was protective against oxidative injury induced by NMDA or transient ischemia^22^.

Noxes and the O_2_**^·^**^-^ that they produce play a fundamental role in vascular inflammation^23,24^ O_2_**^·^**^-^ is the conjugate base of HO_2_**^·^** and is rapidly converted to H_2_O_2_ spontaneously or enzymatically by superoxide dismutases. O_2_**^·^**^-^ can also react with nitric oxide to form peroxynitrite^25^, with metals to create more reactive oxidants (e.g. OH^•^), with membrane phospholipids to form lipid peroxides, or it can directly oxidize proteins^26^. Association of Nox1 with LRRC8A, and localized influx of O_2_**^·^**^-^ may allow selective targeting of signaling proteins that bind to the cytoplasmic leucine rich domains of the channel^21^. In this scenario the short half-life and limited diffusion distance of O_2_**^·^**^-^ places useful limits on off-target oxidation.

Atherosclerosis is a chronic vascular inflammatory disease linked to hypercholesterolemia-induced oxidative stress^27^. It underlies the pathogenesis of coronary artery disease and stroke, the top two causes of death worldwide. VSMCs play a critical pathophysiological role as lineage tracing has revealed that the majority of the fat-containing foam cells in the vascular wall are of smooth muscle origin^28^. Atherosclerotic VSMCs are inflamed^29^, under oxidative stress^30^, have incurred telomere damage^31^, are more frequently senescent^32^, and utilize glycolysis as a preferred energy source^33^. These changes are driven by activation of scavenger receptors such as LOX-1 and CD36 that take up oxidized LDL^34^.

Here we provide direct evidence that VRACs promote O_2_**^·^**^-^ access to the cytoplasm. We then explore how the loss of these channels in VSMCs impacts gene expression, oxidative stress, mitochondrial function, metabolism, proliferation and signaling pathways related to inflammation, senescence and longevity. Compared to wild type (WT) controls, cultured LRRC8A knockout (KO) cells are under less oxidative stress, utilize less energy, and are resistant to stimuli that promote vascular disease. This motivated an assessment of how losing these channels, specifically in VSMCs, affected the development of atherosclerosis and resistance artery dysfunction in hypercholesterolemic mice lacking the ApoE receptor (ApoE^-/-^). Consistent with the *in vitro* characteristics of cultured KO VSMCs, smooth muscle LRRC8A KO mice, and to a significant degree heterozygous animals (HET), were protected from disease.

## METHODS

A detailed description of all materials and methods is provided in the Data Supplement.

### Animal Studies

All animal procedures were approved by the Institutional Animal Care and Use Committee of Vanderbilt University Medical Center and complied with the ARRIVE guidelines. Vascular smooth muscle cell (VSMC)-specific *Lrrc8a* knockout (KO) mice were generated by crossing *Lrrc8a*^fl/fl^ mice with transgenic mice expressing Cre recombinase under the SM22α promoter. To induce atherosclerosis, these mice were crossed onto an Apolipoprotein E-null (*ApoE*^−/−^) background and fed a high-fat diet (42% Fat, TD88137, Harlan Teklad) for 15 weeks after weaning. Aortic atherosclerotic plaque burden was quantified by oil-red-O staining and mesenteric vascular function was assessed using wire myography.

### Cell Culture and *in vitro* Assays

Primary aortic VSMCs were isolated from male KO and WT littermates as previously described^35^ and used for experiments between passages 6 and 15. Cellular redox status was assessed in dye-loaded cells by confocal microscopy (for DHE and ROSstar 550 imaging) and flow cytometry (for DHE, ROSstar 550, MitoSOX, and TMRE), and by measuring superoxide influx, mitochondrial/cytosolic superoxide, membrane lipid peroxidation (MDA assay), and the reduced-to-oxidized glutathione (GSH/GSSG) ratio. Metabolic flux was analyzed using a Seahorse XF Analyzer to determine the extracellular acidification rate (ECAR) and oxygen consumption rate (OCR). VSMC proliferation, migration, and senescence were quantified by MTT, scratch-wound healing assays, and senescence-associated β-galactosidase activity, respectively. Uptake of fluorescently labeled Dil-OxLDL (ThermoFisher) was quantified by confocal microscopy using ImageJ software.

### Molecular and Signaling Analyses

Global transcriptomic changes were profiled by bulk RNA sequencing (Illumina NovaSeq), with differential expression analysis performed in Partek Flow using the DESeq2 algorithm. Key protein expression and phosphorylation status of signaling intermediates (e.g., pEGFR, pAKT) were validated by Western blotting using an infrared imaging system (LI-COR Odyssey).

Transcription factor activity (NF-κB, Hif-1α, Nrf2) was quantified using lentiviral luciferase reporters, with activity normalized for infection efficiency. Relative telomere length and selected mRNA transcripts (e.g., Rps4l, Nrf2 targets, Hk2, Pfkp) were quantified by real-time quantitative PCR (qPCR) as detailed in the Data Supplement. Telomere length was measured using a telomere/single-copy gene (T/S) assay on genomic DNA, whereas gene-expression qPCR was performed on cDNA generated from total RNA, with normalization to housekeeping transcripts.

### Statistical Analysis

Data are presented as mean ± SEM. Statistical analyses were performed using GraphPad Prism 9. Comparisons were made using a one sample *t*- test, unpaired *t*-test or one-way ANOVA with Tukey’s post hoc test, as appropriate. A (\**P*<0.05, \*\**P*<0.01, \*\*\**P*<0.001) was considered statistically significant.

## RESULTS

### Superoxide flux

To detect influx of O_2_**^·^**^-^, unstimulated WT and LRRC8A KO HEK293 cells were loaded with DHE (5 µM) for 30 min, washed 3 times and then exposed to O_2_**^·^**^-^ for an additional 30 min under culture conditions (McCoy’s 5A medium) (Fig. 1A). Oxidized (2-OH)-ethidium, a highly specific reaction product of ethidium and O_2_**^·^**^-^, was quantified by HPLC and normalized to total protein (pmol/mg protein). Cytoplasmic ethidium was oxidized by extracellular O_2_**^·^**^-^ in WT but not LRRC8A KO cells (Fig. 1A). To control the nature and activity of the VRACs present in the plasma membrane, HCT116 cells lacking all LRRC8 isoforms (LRRC8^-/-^) were made to express LRRC8A, or LRRC8C or LRRC8D with the LRRC8A intracellular loop and C-terminal GFP tags. Under isotonic conditions, tagged homomeric LRRC8A channels are minimally active^36,37^. However, when LRRC8C or 8D channels are given the intracellular loop of LRRC8A they are trafficked to the plasma membrane^38^, and incorporation of a C-terminal GFP tag renders them constitutively active under isotonic conditions^16,37^. DHE loaded cells were exposed to extracellular O_2_**^·^**^-^ (Fig. 1B), then fixed and imaged by confocal microscopy (Fig. 1C).

**Figure 1.**
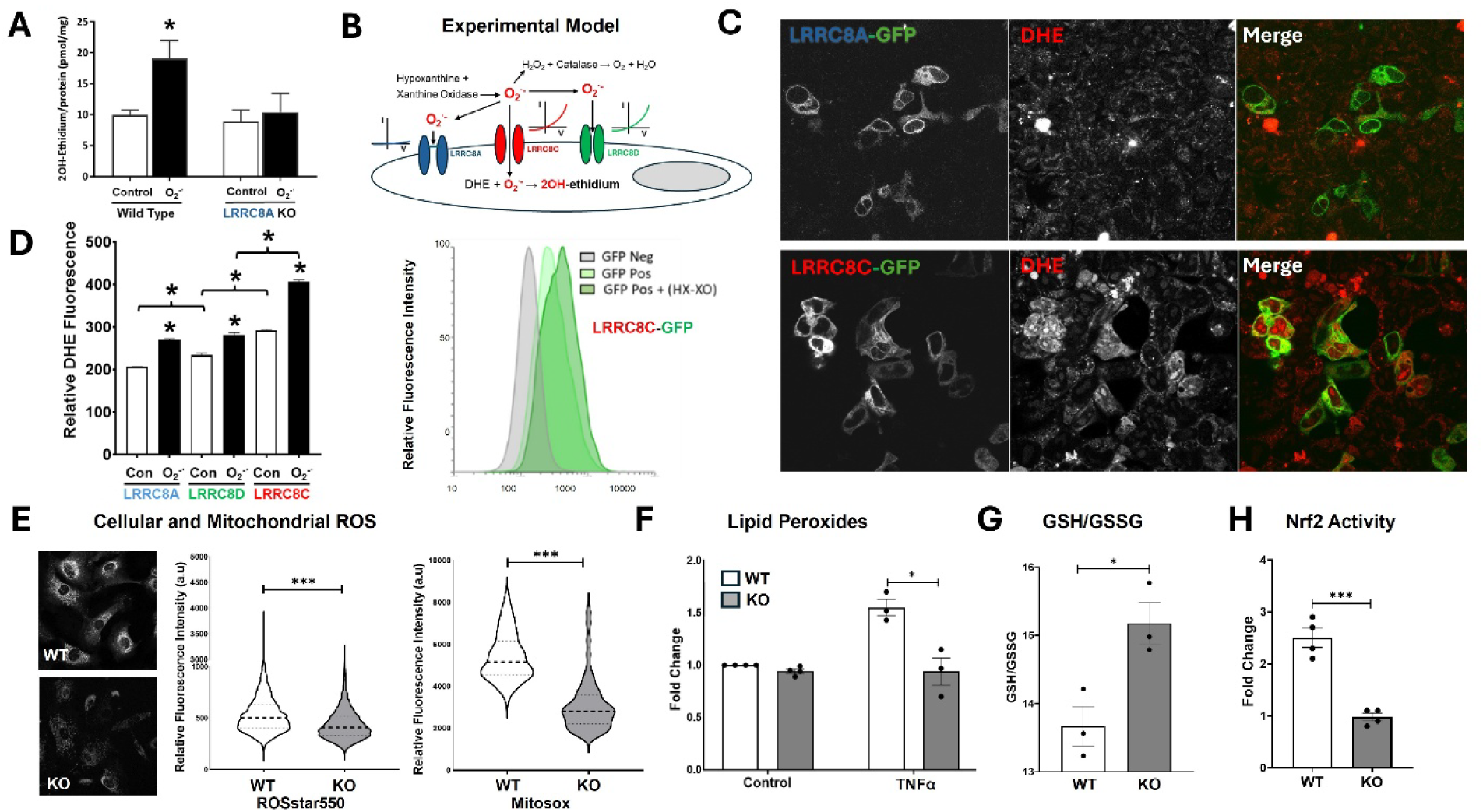
LRRC8 channels support superoxide influx and knockout reduce oxidative stress. A, WT and LRRC8A KO HEK293 cells were loaded with dihydroethidium (DHE), washed, and exposed to extracellular O_2_**^·^**^-^ generated by hypoxanthine/xanthine oxidase (HX–XO). 2-OH-ethidium was quantified by HPLC and normalized to protein. O_2_**^·^**^-^ increased 2-OH-ethidium in WT but not KO cells (mean ± SEM; n = 3 independent experiments; two-way ANOVA). **B,** Protocol for experiments in (**C)** and (**D)**: GFP-tagged LRRC8 proteins were expressed in DHE-loaded HCT116 cells lacking all other LRRC8 proteins and exposed to O_2_**^·^**^-^. **C,** Confocal LRRC8A-GFP (top) or LRRC8C-GFP (bottom); GFP tag (left), DHE (middle) and merged images (right). Cells that express LRRC8C but not LRRC8A have higher DHE fluorescence. **D,** *Left:* Flow-cytometric quantification of DHE fluorescence in LRRC8A-, LRRC8D-, or LRRC8C-GFP–expressing cells under control and O_2_**^·^**^-^-exposed conditions (2-way ANOVA); *Right:* overlaid histograms for LRRC8C-GFP cells demonstrating increased fluorescence in cells that express LRRC8C-GFP and an additional increase in cells exposed to O_2_**^·^**^-^. **E,** ROS imaging in WT and KO primary aortic VSMCs using ROSstar550 or MitoSOX; *Left*: confocal images of cells exposed to ROSstar550. *Right*: Quantification of ROSstar and Mitosox signals by flow cytometry. Violin plots show single-cell fluorescence (dye-positive gate, all cells from 3 independent experiments, Welch’s *t* test). **F,** Lipid peroxidation (malondialdehyde adducts) at baseline and after TNFα. The difference at baseline was not significant (P = 0.07) but TNFα increased peroxidation only in WT cells (for ctrl n = 4 and for Tnf treatment n= 3). **G,** Reduced/oxidized glutathione (GSH/GSSG) under resting conditions; KO had a significantly reduced ratio (n = 3). **H,** Nrf2 reporter activity under resting conditions is lower in KO VSMCs (n = 4). All bars show mean ± SEM and dots represent independent replicates. *P<0.05, and ***P<0.001.

Transfection efficiency was less than 100%, allowing comparison of the DHE signal between expressing (GFP positive) and non-expressing (GFP negative) cells in single microscopic fields.

Expression of LRRC8A-GFP or LRRC8D-GFP had no visually discernable effect on DHE fluorescence in either resting or O_2_**^·^**^-^-exposed cells, while the presence of LRRC8C-GFP cell-specifically increased the subjective DHE signal after O_2_**^·^**^-^ treatment. To quantify this association, flow cytometry was used to compare DHE signal intensity under control conditions and after exposure to O_2_**^·^**^-^ (Fig 1C). Resting LRRC8C-GFP and LRRC8D-GFP expressing cells both had a higher average resting DHE fluorescence than those expressing inactive LRRC8A-GFP, and this difference was larger in cells expressing LRRC8C. Following exposure to O_2_**^·^**^-^ the DHE signal increased in all cells. This increase did not differ between LRRC8A-GFP and LRRC8D-GFP expressors but was significantly greater in cells expressing LRRC8C-GFP. This suggests that LRRC8C channels support O_2_**^·^**^-^ production and influx better than LRRC8A or LRRC8D.

### Oxidative Stress

NADPH oxidase activation can trigger mitochondrial O_2_**^·^**^-^ production in a feed-forward cycle of ROS production^39^. We hypothesized that reduced Nox activity and O_2_**^·^**^-^ influx in LRRC8A KO VSMCs^17^ would reduce mitochondrial ROS production. Confocal microscopy confirmed that ROSstar (cytoplasmic + mitochondria) intensity appeared to be markedly lower in KO cells compared to WT, indicating reduced overall ROS levels. Therefore MitoSOX (mitochondria-specific) and ROSstar oxidant signals were compared by flow cytometry, and both were significantly reduced in KO compared to WT cells (Fig. 1E).

Membrane lipid peroxidation occurs under oxidant stress. This was quantified under resting conditions and after TNFα (10 ng/ml) exposure. Resting levels were lower in KO cells but not significantly so (P = 0.06, N = 4) but TNFα stimulation increased MDA levels only in WT cells (Fig. 1F). The ratio of reduced to oxidized glutathione (GSH/GSSG) was significantly higher in KO cells (Fig. 1G), indicative of a more reduced cytoplasmic environment. As an integrated assessment of oxidative stress, Nrf2 transcriptional activity was measured using a luciferase reporter. Nrf2 activity was significantly lower in KO VSMCs (Fig. 1H), consistent with a reduced need for antioxidant protection. Protein products of Nrf2 target genes, including extracellular superoxide dismutase (SOD3) and heme oxygenase (HO-1) were reduced in KO cells (Suppl. Fig. 1A), as was the expression of mRNA (qPCR) for other Nrf2 target genes including glutathione peroxidase 3 (*Gpx3*), glutathione S-transferase alpha 3 (*Gsta3*), and heme oxygenase-1 (*Hmox1*) (Suppl. Fig. 1B). The protein abundance of the membrane components of the Nox1 complex (Nox1, p22phox) were unchanged in LRRC8A KO VSMCs. However, the level of the cytosolic activator Noxa1 subunit was significantly lower in KO cells. Of note, VSMC-specific KO of Noxa1 protected against both neointimal hyperplasia following endovascular injury and atheroma formation in ApoE^-/-^ mice^40^. These data, and many others were normalized to tubulin which was not differentially expressed in KO cells (Suppl. Fig. 1D).

### Transcriptomic Analysis and HIF-1α Signaling

Bulk RNA sequencing was performed on primary cultured aortic VSMCs from 3 pairs of WT and KO littermates (Suppl. Fig. 2A). Principal component analysis showed clear segregation of WT and KO samples and a heatmap of global gene expression patterns indicated distinct clustering by genotype (Suppl. Fig. 2 B and C). Differential expression analysis using DESeq2 identified 573 differentially expressed genes (DEGs) between KO and WT VSMCs. Among these, 206 genes were upregulated, and 367 genes downregulated in KO cells (p ≤ 0.05 and Fold Change ≥ 1.5). A volcano plot is provided in Suppl. Fig. 2D.

To interpret the impact of these transcriptional changes we performed pathway enrichment analysis using KEGG pathways via Enrichr. Fig. 2A presents the top eight affected KEGG pathways based on all DEGs. Genes showing increased expression are in red and decreased genes are in green with bar height indicating the combined score/enrichment. Altered genes were most associated with inflammation (Hif-1α, AGE-RAGE, Hippo and MAPK signaling), and metabolism (lipolysis, glutathione metabolism, Rap1 signaling. When analysis was confined to the 367 downregulated genes the most affected functional categories were the Hif-1α pathway, stress-related metabolic processes (glutathione metabolism, glycolysis), and inflammatory signaling (Fig. 2B, right panel). Numerous NF-κB target genes and cytokine signaling components were expressed at significantly lower levels in KO VSMCs at baseline, including receptors for interleukin-6 (Il6ra) and interleukin-1β (Il1r1). Read counts for strongly repressed genes in key inflammatory pathways are provided in Suppl. Fig. 3. Analysis of upregulated transcripts is shown in Suppl. Fig. 4 shows enrichment of Rap1 signaling. This small GTPase regulates proliferation and migration of VSMCs^41^. Collectively, the RNAseq data indicate that loss of LRRC8A induces an anti-inflammatory transcriptional profile that is coupled to significant alterations in metabolism and stress-related processes. Accordingly, an analysis of gene-disease association for the downregulated gene set using the DisGeNET database revealed a link to vascular inflammation associated with atherosclerosis and myocardial infarction (Suppl. Fig. 2E).

**Figure 2.**
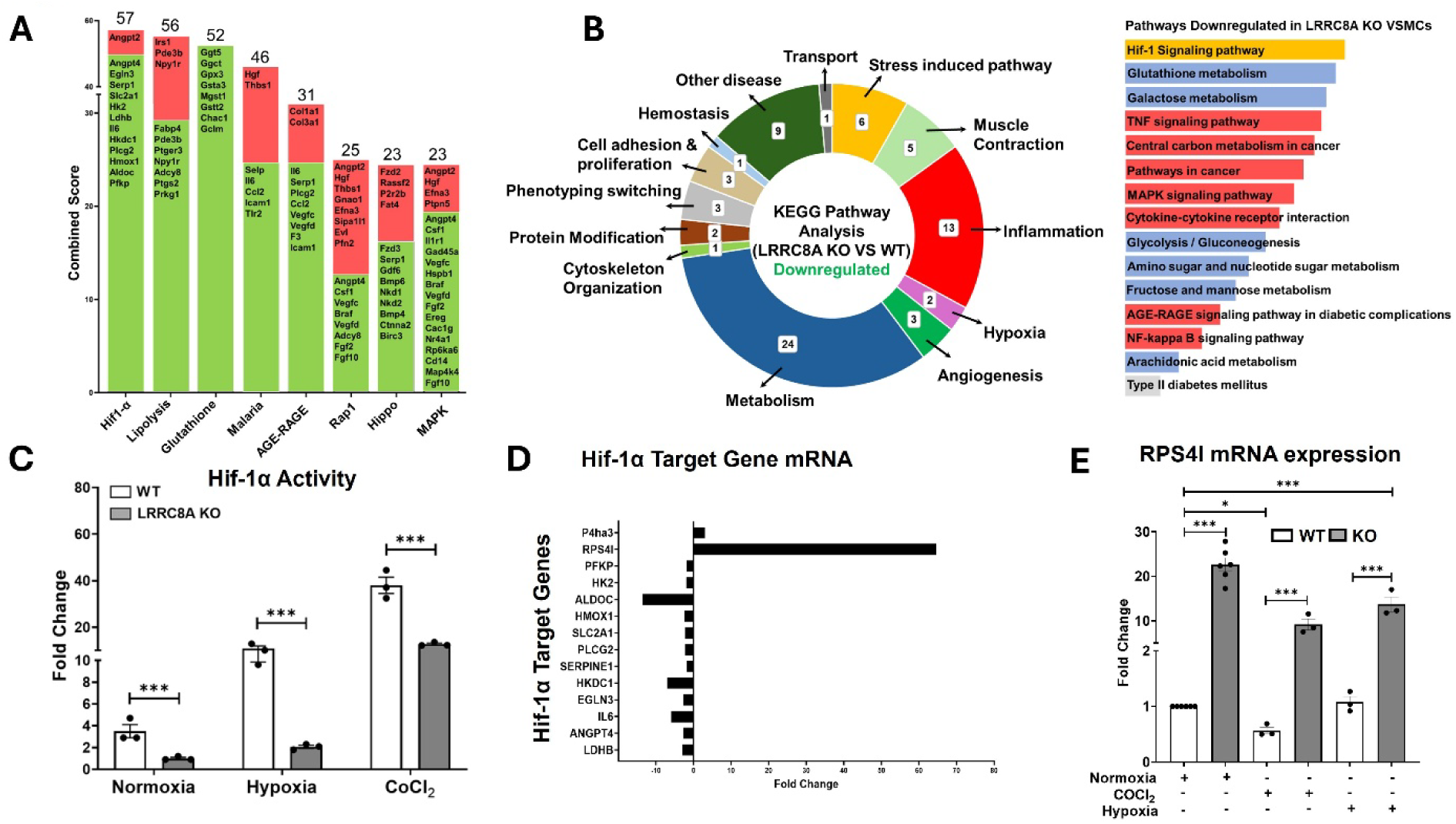
Effects of LRRC8A KO on mRNA expression and Hif1α signaling in VSMCs. A, Most altered KEGG pathways (combined score) with increased (red) and decreased (green) genes noted. **B,** Categories of downregulated genes and associated pathways. **C.** Hif-1α reporter (HRE-luciferase) activity in WT and KO VSMCs under normoxia, hypoxia (1% O₂, 24 h), or CoCl₂ (300 µM, 6 h) stimulation; activity normalized to protein and corrected for reporter infection efficiency (GFP/tubulin). (normoxia, one-sample *t* test; hypoxia and CoCl₂, Mann–Whitney test; n = 3). **D,** RNA-seq log₂ fold change (KO vs WT) for canonical HIF-1α target genes. **E,** qPCR validation of Rps4l expression under the same conditions as in (**C)** normalized to WT in normoxia (normoxia, one-sample *t* test; hypoxia and CoCl₂, Mann–Whitney test; n = 6 for control and n = 3 for hypoxia and CoCl₂). Bars are mean ± SEM, dots represent replicates. *P<0.05, ***P<0.001.

As Hif-1α signaling was the most significantly altered pathway this was assessed directly using a hypoxia response element (HRE)-luciferase reporter. Activity was ∼3X higher in resting WT cells. Hypoxia (1% O₂, 24 h) and chemical induction with CoCl₂ induced a similar fold change compared to baseline, but WT remained significantly greater than KO under all conditions (Fig. 2C). Analysis of the expression of a panel of canonical Hif-1α target genes revealed consistent downregulation in KO cells (Fig. 2D). A clear exception was Rps4l, a known target of HIF-1α-mediated repression, with a read count that was elevated over 60-fold in KO VSMCs (Fig. 2D). This gene has both long-noncoding and peptide-encoding functions, both of which impair S6 kinase activity and reduce VSMC proliferation and VSMC Rps4l expression is protective in pulmonary hypertension^42,43^. To further explore the regulation of Rps4l qPCR was performed under control conditions and after exposure to hypoxia or CoCl_2_. Expression decreased in response to both stimuli but remained markedly higher in KO cells under all conditions (Fig. 2E). Hif-1α is a critical regulator of glycolysis. Under control conditions expression of 2 hexokinase (Hk2) was reduced in KO cells, while that of phosphofructokinase (Pfkp) was not. Both genes displayed impaired transcriptional responses to hypoxia in KO cells (Suppl. Fig. 5).

### Metabolism, Proliferation and Migration

Transcriptomic analysis suggested downregulation of glycolysis in KO cells. This included decreased expression of Glut1 (Slc2a1, 2.3X, p < 0.0001) which mediates insulin-independent glucose uptake. Insulin-sensitive Glut4 was not detected. We assessed the abundance of Hk2, pyruvate kinase M2 (Pkm), and 6-phosphofructo-2-kinase/fructose-2,6-biphosphatase-3 (Pfkfb3) protein and found significant decreases in all three. The latter is a master regulator of glycolytic flux in VSMCs and is upregulated in intimal hyperplasia^44^ (Fig. 3A). We then directly assessed the metabolic activity of WT and KO VSMCs using an XF extracellular flux analyzer.

**Figure 3.**
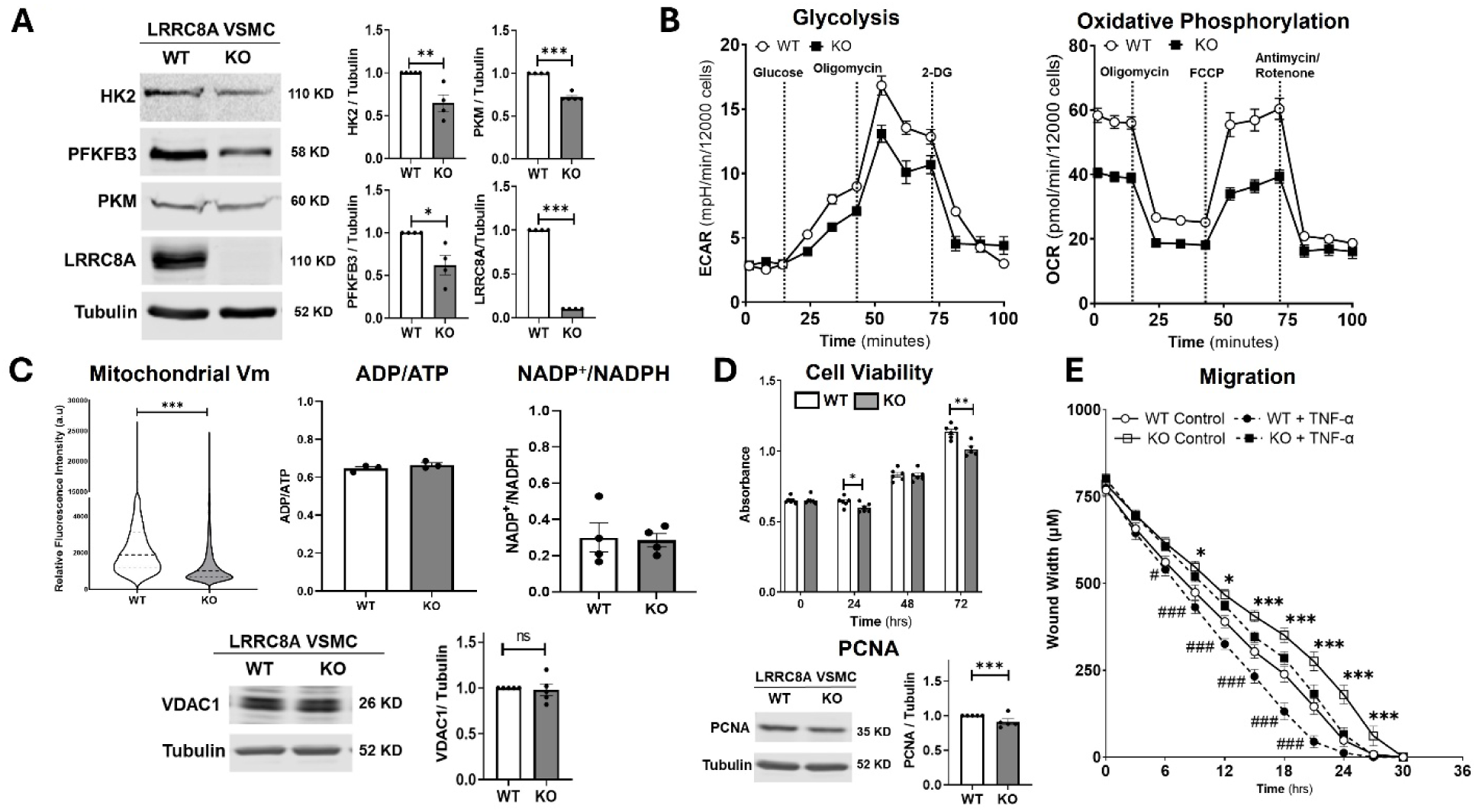
Metabolism and energy balance in LRRC8A KO VSMCs. A, Abundance of HK2, PFKFB3, PKM, and LRRC8A in primary aortic VSMCs (WT, KO) normalized to α-tubulin (one sample *t* test; *n* = 4 independent lysates per genotype). **B,** Seahorse glycolytic stress test (ECAR; injections: glucose, oligomycin, 2-deoxyglucose) and mitochondrial stress test (OCR; injections: oligomycin, FCCP, antimycin/rotenone) n = 4. **C,** Mitochondrial membrane potential comparison by TMRE using flow cytometry (violin plots of single-cell fluorescence, dye-positive gate) (pooled cells from *n* = 3 independent experiments, unpaired *t* test). ADP/ATP (*n* = 3) and NADP⁺/NADPH (*n* = 4), Welch’s *t* test and VDAC1 abundance as an estimate of mitochondrial biomass. (n = 5, one sample *t* test). **D,** Cell viability assays using MTT over 72 h (n = 6, One-way ANOVA), and PCNA immunoblotting. (unpaired *t* test; *n* = 4). **E,** Migration assessed by scratch-wound assay (IncuCyte) under control conditions and after TNFα stimulation (10 ng/mL). Wound width (µm) was monitored over 72 h; two-way ANOVA was used to compare WT vs KO under control and TNFα conditions (n = 3 biological replicates with 6 wells per condition). Bars/plots show mean ± SEM; dots represent independent replicates. *P<0.05, ***P<0.001 for WT vs KO; #P<0.05, ###P<0.001 for TNFα vs control within genotype.

Extracellular acidification rate (ECAR) revealed a significantly lower glucose-stimulated glycolytic rate in KO cells but no difference in maximal glycolytic capacity or glycolytic reserve (Fig. 3B, left panel; Suppl. Fig. 6, left panel).

Oxidative phosphorylation (OCR) was also suppressed in KO cells. Significant reductions were observed in all major parameters, including basal respiration, ATP-linked respiration, maximal respiratory capacity and non-mitochondrial respiration (Fig. 3B, right panel; Suppl. Fig. 6, right panel). This was not associated with a change in proton leak, indicating that ATP production was not differentially uncoupled. OCR was previously reported to be reduced in skeletal muscle myotubes lacking LRRC8A^45^. Reduced OCR was associated with a reduction in mitochondrial membrane potential (ΔΨm) as assessed by TMRE staining (Fig. 3C, left panel), indicating a reduced proton gradient. VDAC1 abundance correlates with mitochondrial mass and was not altered in KO cells (Fig. 3C, bottom panel). To discern if a reduced ΔΨm reflects mitochondrial dysfunction or lower energy demand we assessed the availability of ATP and reducing equivalents as reflected by ADP/ATP and NADP^+^/NADPH respectively, both were unaltered (Fig. 3C, middle and right panel). Maintenance of energy homeostasis in the setting of lower energy production points to lower demand. Proliferation increases demand and a proliferative VSMC phenotype is associated with upregulation of glycolysis. MTT assays demonstrated a decrease in proliferation that was confirmed a reduced abundance of Proliferating Cell Nuclear Antigen (PCNA) in KO cells (Fig. 3D). Movement also required significant energy and the migratory rate of KO cells was reduced both at rest and following TNFα stimulation in a scratch assay (Fig. 3E). Collectively, these results indicate that loss of LRRC8A in VSMCs reduced metabolic rate.

### Epidermal Growth Factor Signaling

Protein-Protein Interaction (PPI) hub analysis of the RNAseq data identified significantly enriched pathways as visualized by heatmap clustering (Figure 4A). The top hits were two receptor tyrosine kinases, the epidermal growth factor receptor (EGFR) and the type 1 insulin-like growth factor receptor (IGF1R). Differentially expressed genes associated with the EGFR hub included key adaptor and regulatory proteins (e.g., Grb10, Sh3kbp1, Ptpre, Socs3), multiple ligands and growth factors (e.g., Hbegf, Ereg, Hgf, Vegfc), cell adhesion molecules (e.g., Icam1, Ncam1, Tnc) and downstream signaling effectors (Fig. 4A). We then assessed the functional status of EGFR signaling. Under resting conditions, the abundance of EGFR protein was unchanged but phosphorylation at Y1068 (pEGFR), an autophosphorylation site that reflects activation, was significantly reduced in KO cells (Fig. 4B). Insulin, EGFR and IGF1R all activate the PI3K/AKT cascade to promote glycolysis, cell growth and survival, and to inhibit lipolysis (the second most altered pathway based on RNAseq (Fig. 2A). EGFR activates PI3 kinase to create phosphinositol (3,4,5)-triphosphate (PIP3) which coordinates PIP3-Dependent Kinase 1 (PDK1), Mechanistic Target of Rapamycin Complex 2 (mTORC2), and AKT at the plasma membrane. PDK1 and mTORC2 phosphorylate AKT at T308 and S473 respectively to cause activation. There was a significant increase in total AKT protein in KO cells but pAKT was reduced at both sites. Exposure to EGF (50ng/ml) increased pEGFR but the difference between WT and KO only remained statistically significant at T308 pointing to greater impact on PDK1 (Fig. 4C). In view of the LRRC8A-dependence of AKT activation by insulin and the shared utilization of Insulin Receptor Substrate 1 (IRS1) by the insulin and EGF receptors we assessed expression. Western blotting confirmed what was suggested by RNAseq, a significant increase in IRS1 protein in KO cells (Fig. 4D).

**Figure 4.**
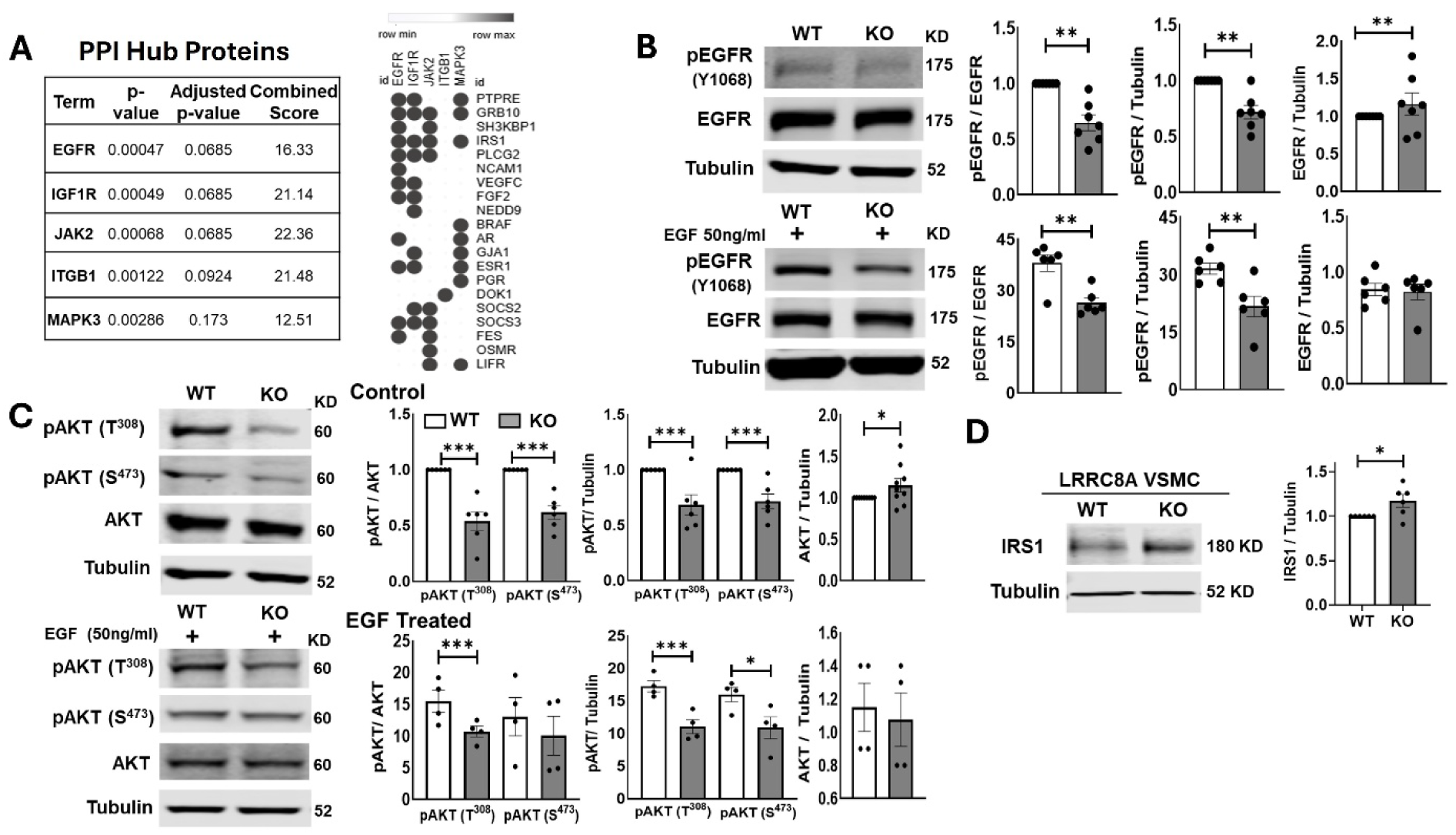
EGFR-centered network and EGF–AKT signaling in LRRC8A-deficient VSMCs. A, Top-ranked Protein–Protein Interaction (PPI) hubs from RNAseq analysis (KO vs WT) with associated partner proteins shown in dot map; EGFR and IGF1R were the most significantly enriched hubs. B, Immunoblotting for phosphorylated EGFR (pY1068) and total EGFR under basal conditions and after EGF stimulation (50 ng/mL, 10 min); signals normalized to total EGFR or α-tubulin. For control lanes, n = 7, one-sample *t* test; for EGF-treated comparisons, n = 6, unpaired *t* test. **C,** Immunoblotting for phosphorylated AKT (pT308, pS473) and total AKT under basal conditions and after EGF stimulation; normalized to total AKT or α-tubulin. For control, n = 6, one-sample *t* test; for EGF-treated WT vs KO, n = 4, unpaired *t* test. D, IRS1 protein abundance normalized to α-tubulin (n = 6, one sample *t* test). Bars and plots show mean ± SEM with individual biological replicates; significance indicated on panels (*P<0.05, **P<0.01, ***P<0.001).

### Inflammation and Senescence

RNAseq pointed to altered TNFα/NF-κB signaling, as we reported previously following siRNA knockdown of LRRC8A or 8C^16,17,29^. We used an NF-κB-luciferase reporter to confirm that both basal and TNFα-stimulated NF-κB activity were reduced >10-fold in KO compared to WT VSMCs (Fig. 5A). However, KO cells remained responsive to TNFα based on fold change from this much lower baseline level.

**Figure 5.**
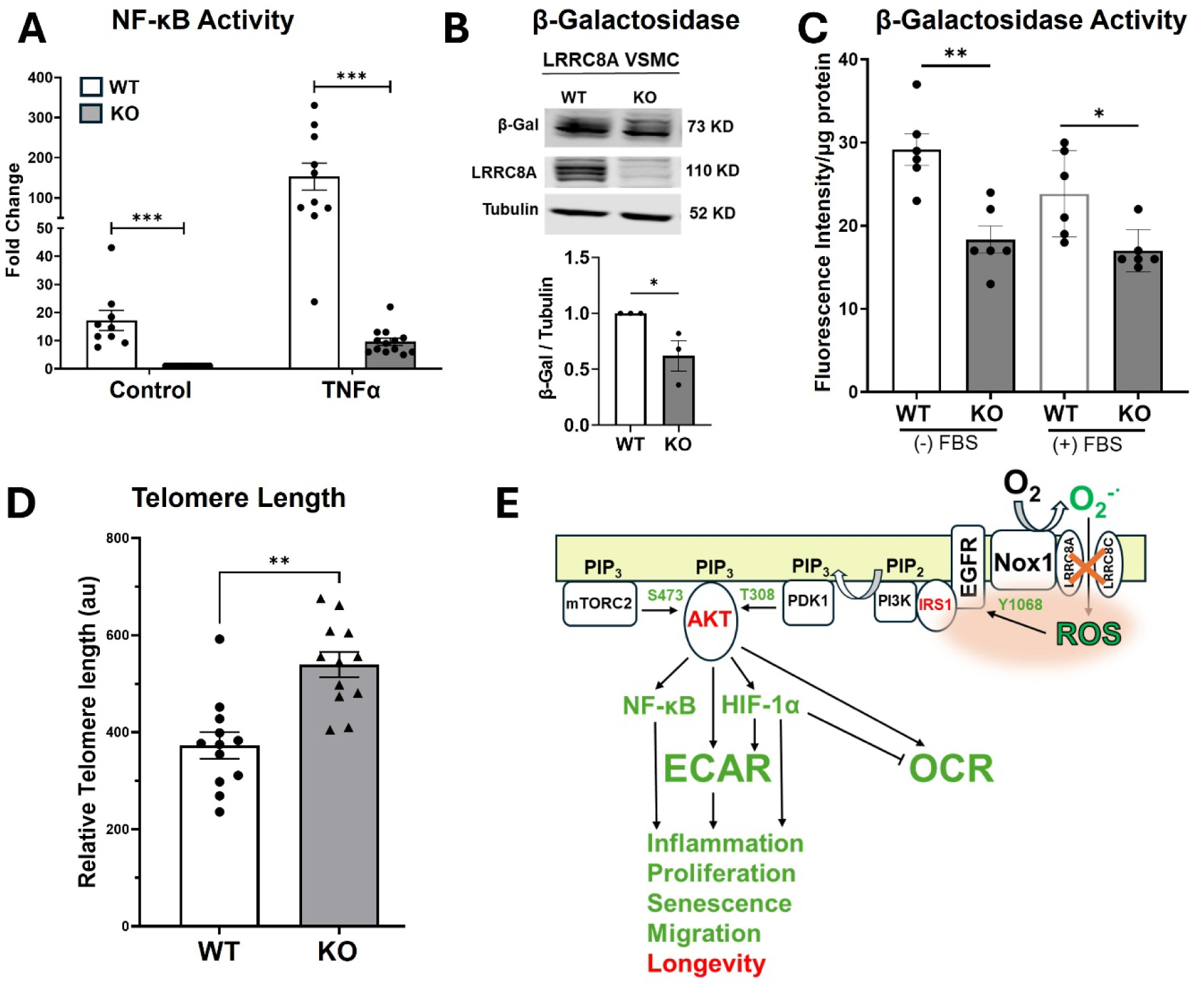
NF-κB Activity, Senescence Markers, Telomere Length and Mechanistic Model of LRRC8A Function in VSMCs. A, NF-κB reporter activity in WT and KO VSMCs under control conditions and after TNFα (10 ng/mL, 6 h) normalized to protein and corrected for infection efficiency (GFP/tubulin) and expressed as fold change from control KO cells, (control samples, one-sample *t* test; for TNF treated samples Mann-Whitney test; n ≥ 9). **B,** β-galactosidase (β-Gal) protein abundance in WT and KO VSMCs normalized to α-tubulin (n = 3, unpaired *t* test). **C,** β-Gal enzymatic activity (fluorescence/µg protein) in WT and KO VSMCs grown in either low (*left*) or normal (*right*) serum (n = 6, One-way ANOVA). **D,** Relative telomere length (qPCR T/S ratio, arbitrary units) in WT and KO VSMCs (n = 12, unpaired *t* test). **E,** Schematic model linking Nox1/LRRC8A/O **^·^**^-^ and EGFR/AKT signaling to NF-κB and HIF-1α pathways and effects on ECAR/OCR, proliferation, migration, senescence, and longevity. Green text indicates pathways or processes that are decreased in LRRC8A KO VSMCs, and red text indicates those that are increased. Means ± SEM. \**P*<0.05, \*\**P<0.01,* \*\*\**P<0.001*.

Reduced oxidative stress, a low metabolic rate, and less inflammation should have significant phenotypic consequences. We assessed mRNA expression for known markers of contractile vs. secretory VSMCs (Suppl. Table 1). There was a significant reduction in 3 of 23 markers of a contractile phenotype (*Itga1, Myh11, Lmod1*), and no consistent upregulation of synthetic markers (*Col1a1* up, *Ereg* down). There were no indications of phenotypic transformation (fibroblast, fat, bone, etc.), nor changes in autophagy or apoptosis markers. However, there was decreased expression of 38% (15/39) of established markers of senescence including cell cycle inhibitors (*Cdkn1a, Cdkn2b*) and the lysosomal enzyme β-galactosidase (β-Gal, encoded by *Glb1*). Western blot analysis showed reduced β-Gal protein, which correlated with lower SA-β-galactosidase activity in KO cells (Fig. 5B and C). Sustained stress drives senescence partly through telomere attrition. LRRC8A KO VSMCs had significantly longer telomeres than WT (Fig. 5D), suggestive of enhanced genomic stability and greater replicative capacity.

Changes in cultured VSMCs are summarized in Fig. 5E. Loss of LRRC8A impairs redox-dependent activation of the EGFR, reducing PIP3 production and AKT activation. This impairs Hif-1α and NF-κB-driven inflammation and reduces senescence. This anti-inflammatory and anti-senescent phenotype in cultured cells provided the impetus to assess the impact of VSMC-specific LRRC8A KO in hypercholesterolemia *in vivo*.

### Atherosclerosis

WT and VSMC-specific LRRC8A knockout (KO) mice were bred into an ApoE^-/-^background to create hypercholesterolemic mice with 15 weeks on a high fat diet. Serum cholesterol and triglycerides did not differ between WT, HET (LRRC8A^+/-^), and KO mice (Suppl. Fig. 7A and B). Oil Red-O staining was performed on aortae to visualize atherosclerotic plaque coverage. LRRC8A KO and HET aortae from both male and female mice showed a significant reduction in lesion area compared to WT (Fig. 6A), and HET lesion area was not significantly different from KO.

**Figure 6.**
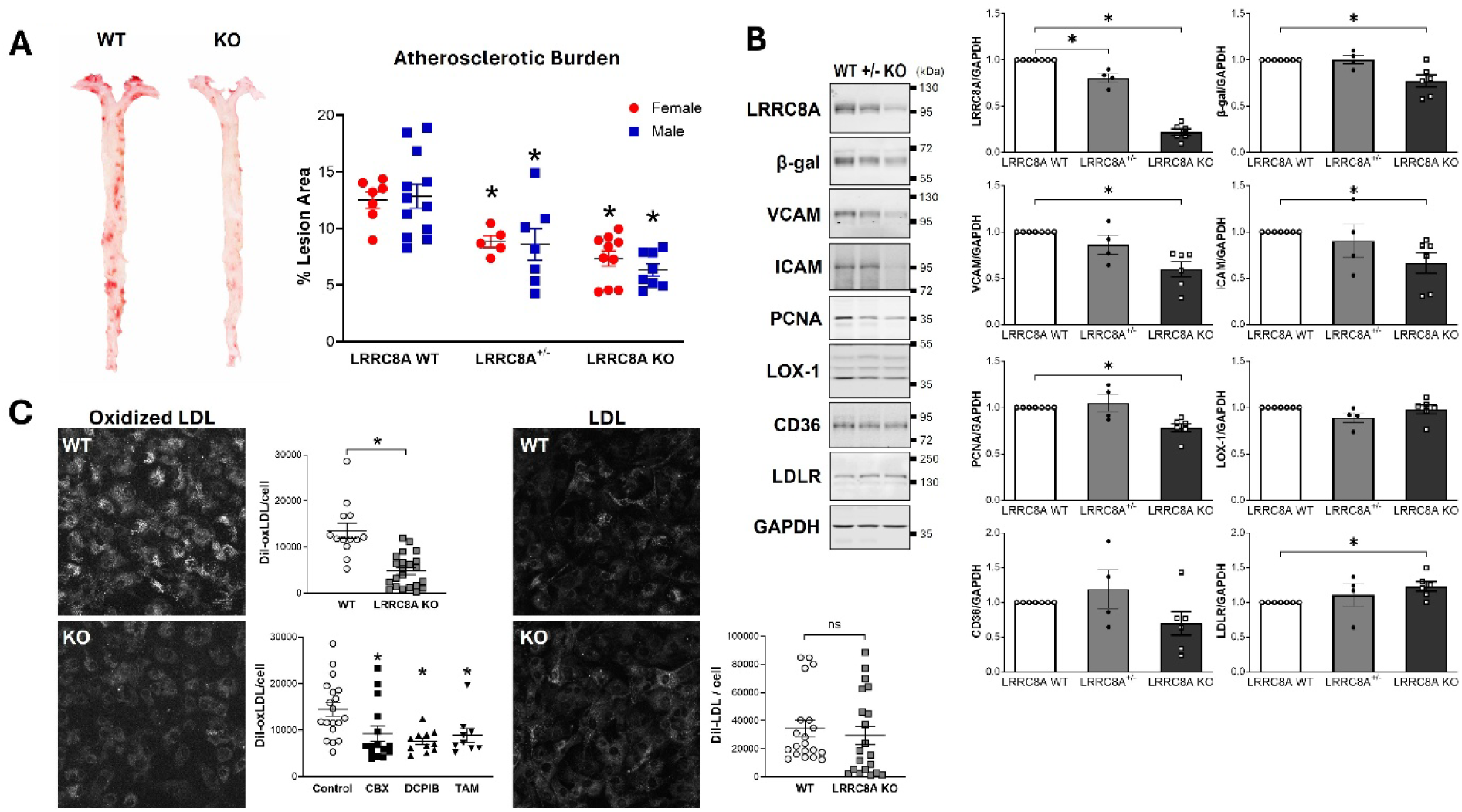
Reduced aortic atherosclerosis, inflammation, proliferation and senescence in VSMC-specific LRRC8A KO mice are associated with impaired oxidized-LDL uptake in VSMC. A, Aortic atherosclerotic lesions. Representative en-face Oil Red O–stained aortae from a WT and a KO mouse, and quantification of percent lipid-stained area relative to total aortic area in male and female ApoE^-/-^;SM22α-Cre LRRC8A genotypes (WT, +/-, KO) mice (Mann-Whitney test). \**p* < 0.05 compared to WT in each sex. **B,** Aortic protein expression. Immunoblotting of aortic lysates from male ApoE^-/-^, LRRC8A WT, HET (+/-) or KO mice normalized to GAPDH for LRRC8A, β-Gal, VCAM, ICAM, PCNA, LOX-1, CD36, and LDLR with densitometric quantification. Each point represents one animal. Bars are mean ± SEM. \**p* < 0.05. **C,** Lipoprotein uptake in primary VSMCs. Confocal quantification of reduced DiI-oxLDL uptake (10 µg/mL, 4 h) in KO VSMCs, or with pharmacologic VRAC inhibition (10 min prior to Dil-oxLDL) in WT cells using carbenoxolone (CBX; 50 µM), DCPIB (30 µM), or tamoxifen (TAM; 10 µM). Uptake of non-oxidized DiI-LDL was not altered in KO cells (Mann-Whitney test). \**p* < 0.05 compared to Control.

LRRC8A protein was dramatically reduced in aortae from KO mice but remained detectable due to protein from non-VSMC cells in the vascular wall. LRRC8A in aortae from male HET mice was intermediate between WT and KO, and significantly lower than WT. A similar pattern was observed for markers of vascular inflammation (ICAM, VCAM), senescence (β-Gal), and proliferation (PCNA) but these changes were only statistically significant in KO vessels (Fig. 6B).

Uptake of oxidized LDL (OxLDL) occurs through scavenger receptors including LOX-1 and CD36 and both pathways trigger ROS production by NADPH oxidases^46,47^. The abundance of LOX-1 and CD36 are increased by their own pro-inflammatory activity^45,46^ but did not differ between WT and KO aortae, while the expression of LDLR was increased in KO aortae from male mice (Fig. 6B). To assess the function of these receptors we compared uptake of Dil-labelled OxLDL (10 µg/ml, 4 hrs) using confocal microscopy. Uptake was reduced in KO compared to WT cells (Fig. 6C). To test the role of VRAC channel function, the impact of three non-structurally related VRAC inhibitors (Carbenoxolone; CBX, DCPIB, Tamoxifen; TAM) on OxLDL uptake by was assessed in WT cells. All three significantly attenuated uptake to a degree that was similar to KO cells. In contrast, uptake of non-oxidized LDL occurs via the LDL receptor (LDLR) which does not activate ROS production. Accumulation of Dil-labeled LDL did not differ between WT and KO cells (Fig. 6C).

### Vascular function

Atherosclerosis enhances vasoconstriction^48^ and impairs both endothelium-dependent and-independent relaxation^49^. Mesenteric arterial vasomotor function in ApoE^-/-^ mice was assessed via wire myography. In LRRC8A WT male and female animals, maximal contraction in response to both norepinephrine (NE, mixed α1-, α2- and β2-adrenergic agonist) and phenylephrine (selective α1) were enhanced compared to HET (LRRC8A^+/-^) or KO mice (Fig. 7A). Endothelium-dependent relaxation to acetylcholine (ACh) was better in female but not male LRRC8A KO and HET rings compared to WT. Endothelium-independent relaxation to sodium nitroprusside (SNP) was significantly improved only in male LRRC8A KO rings.

**Figure 7.**
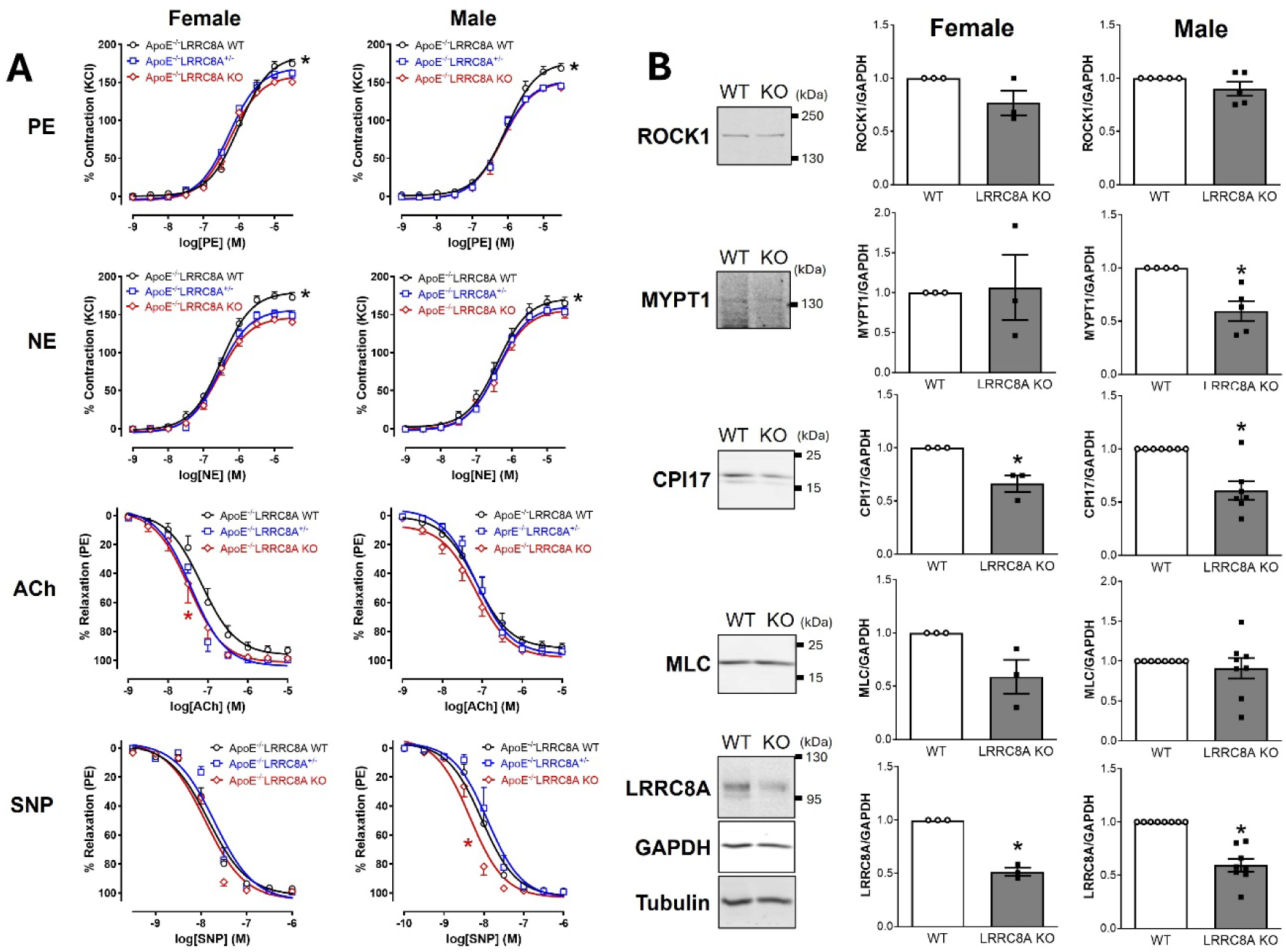
Mesenteric vasomotor function in ApoE^-/-^ mice lacking VSMC LRRC8A. A, Cumulative concentration-response curves for contraction to phenylephrine (PE) and norepinephrine (NE), and relaxation to acetylcholine (ACh, endothelium-dependent) and sodium nitroprusside (SNP, endothelium-independent) in mesenteric arteries from female and male LRRC8A WT, HET (+/-) or KO, ApoE^-/-^ mice. Black asterisks (*) indicate comparisons of maximal responses between WT vs HET or KO. Red asterisks (*) indicate EC_50_ comparisons (ACh: WT vs HET or KO; SNP: WT or HET vs KO, male n = 6-7, female n = 5-6). **B,** Representative immunoblots from isolated mesenteric arteries (WT or KO, ApoE^-/-^) normalized to GAPDH. Bars are mean ± SEM normalized to WT. \**p* < 0.05 (male n = 4 - 8, female n = 3)

Mesenteric artery injury induced by a prolonged (2 day) *in vitro* exposure to TNFα was mitigated by loss of LRRC8A and enhanced relaxation was shown to be associated with reduced RhoA and Rho kinase (ROCK) activation^21^. ROCK phosphorylates myosin phosphatase targeting subunit 1 (MYPT1), the regulatory subunit of myosin light chain phosphatase (MLCP) causing inhibition which promotes vasoconstriction and impairs vasorelaxation. CPI17 can be activated by protein kinase C or ROCK and also inhibits MLCP^50,51^. Western blot analysis of mesenteric artery beds from LRRC8A WT and KO, ApoE^-/-^ mice showed no change in the abundance of ROCK, but MYPT1 expression was significantly decreased in male KO mesenteries. CPI17 protein expression was reduced in KO mice of both sexes compared to WT, while expression of MLC was not altered. Downregulation of MYPT1 and CPI17 in KO vessels may contribute to decreased vasoconstriction and enhanced relaxation. Thus, both plaque progression and vascular dysfunction in ApoE^-/-^ mice is LRRC8A-dependent.

## DISCUSSION

LRRC8 VRACs support Nox1 activity^16,17^. We now show that the function of VRACs affects oxidation of cytoplasmic DHE by extracellular O_2_**^·^**^-^, consistent with our prior proposal that these anion channels support influx of O_2_**^·^**^-^. Experiments defining endpoints linked to redox signaling/stress were guided by changes in gene expression in KO cells. We observed an increase in GSH/GSSG, and reduced mitochondrial O_2_**^·^**^-^ production, lipid peroxidation, Nrf2 activity and expression of antioxidant enzymes. Gene expression reflected decreased inflammatory signaling (NF-κB, Hif1α) and reduced glycolysis. KO cells produced less ATP via both ECAR and OCR and mitochondrial membrane potential was reduced, but neither ATP nor NADPH availability were altered. These changes, in the setting of lower EGFR and AKT activity, reductions in proliferation, motility and senescence, and longer telomeres collectively point to low energy demand in KO VSMCs. We hypothesized that these changes would alter vascular inflammation and oxidative stress in response to hypercholesterolemia. Accordingly, VSMC-specific LRRC8A KO mice were protected from both atheroma formation and increased contractility in an ApoE^-/-^ environment. Oxidized but not reduced LDL uptake was impaired in KO cells, and LRRC8A channel blockers impaired OxLDL uptake in WT cells to a similar degree as seen in LRRC8A KO cells, suggesting for the first time that LRRC8A channels and ROS support endocytosis of OxLDL.

### Superoxide flux and oxidative stress

The ability of extracellular SOD3 to modulate Nox1-dependent activation of intracellular ASK1^19^, and LRRC8A-dependence of MPRIP oxidation^21^ suggested that extracellular O_2_**^·^**^-^ might reach the cytoplasm via these channels^52^. We proposed that influx is promoted by local membrane depolarization produced by current flow through physically-associated Nox1 protein^19,29^. Here we sought to directly link O_2_**^·^**^-^ flux to VRAC activity. Unfortunately, influx of O_2_**^·^**^-^ produced by Nox1 activation cannot be compared because loss or inhibition of VRACs reduces Nox1 activity^17^, and Nox1 activation also increases mitochondrial O_2_**^·^**^-^ production^53,54^. Therefore, we utilized a controlled enzymatic source of extracellular O_2_**^·^**^-^. Unstimulated WT HEK293 cells took up more O_2_**^·^**^-^ than LRRC8A KO HEK293 cells, even though endogenous VRAC activity is low in this setting. We therefore heterologously expressed LRRC8 homomers of known, variable activity^16^. Constitutively active LRRC8C homomers increased the oxidation of cytoplasmic DHE in O_2_**^·^**^-^-exposed cells significantly more than did expression of inactive LRRC8A, or active LRRC8D homomers. This supports the concept that Nox1-dependent redox signaling is best supported by LRRC8A/C channels^16^.

Within the Nox1 complex, the abundance of the integral membrane components Nox1 and p22phox were unaltered in LRRC8A KO cells, but cytoplasmic Noxa1 was reduced. This may reflect reduced inflammation as Noxa1 expression is induced by TNFα and increases in ApoE^-/-^ vessels^55^. Nox1-dependent O_2_**^·^**^-^ can also amplify ROS production by mitochondria via so-called “ROS-induced ROS production”^56^, the mechanism for which is poorly understood. Thus, reduced Nox1 activity and impaired O_2_**^·^**^-^ influx may limit mitochondrial O_2_**^·^**^-^ production in KO cells (Fig. 3C). The combination of these changes is consistent with observed decreases in membrane lipid peroxidation, an increased GHS/GSSG ratio, reduced Nrf2 activity, and altered expression of SOD3 and HO-1.

### Gene Expression

The dominant categories of altered gene expression were metabolism and inflammation, and the most downregulated pathway was Hif1-α which senses hypoxia and drives a metabolic switch to glycolysis. The inflammation associated with atherosclerosis^57^ and pulmonary hypertension^58^ activates Hif-1α and drives aerobic glycolysis (Warburg effect). Hif-1α can also be activated by growth factors and cytokines via PI3K/AKT/mTOR and Ras/MAPK signaling, both of which are triggered by EGFR activation, and impaired by loss of LRRC8A. Hif-1α activity can be modified by multiple factors^59^ including FOXO3a^60^ which interferes with its transcriptional activity. However, FOXO3a activity was impaired in LRRC8A KO VSMCs^61^ making this unlikely to account for reduced Hif-1α activity. RPS4l is a long non-coding RNA that also encodes a peptide, and both mechanisms interfere with RPS6 function to impair proliferation. RPS4l is repressed by Hif-1α and consistent with this was over-expressed >20X in KO cells. This may be of physiological significance as knockdown of RPSl4 in VSMCs promoted pulmonary vascular inflammation and pulmonary hypertension, while overexpression was vasculoprotective^42,43^.

Glutathione provides electrons to reduce oxidized proteins, and under oxidative stress regeneration of reduced glutathione (GSH) requires significant energy to produce NADPH via the pentose phosphate shunt^62^. Thus, reduced oxidative stress lowers energy demand. Decreased expression of GPX3 (Suppl. Fig. 3), a secreted enzyme that acts extracellularly, suggests that extracellular oxidant stress is lower in KO VSMCs. Rap1 signaling was the top pathway identified by analysis of upregulated genes (Suppl. Fig. 4). This small GTPase promotes proliferation and migration of VSMCs^41^ but mice lacking Rap1b were hypertensive, with increased vascular responsiveness to multiple vasoconstrictors^63^. Rap1 regulates ERK signaling^41^ which is impaired by SOD3 and by siLRRC8A^17^, therefore increases in gene expression related to Rap1 signaling may be compensatory.

### Metabolism and signaling

A metabolic shift to aerobic glycolysis (Warburg effect) is associated with vascular injury and atherosclerosis and may drive phenotypic switching^28,33^. An important clue to the protected metabolic state of LRRC8A KO VSMCs was reduced expression of Glut1, suggesting lower demand for glucose uptake. Key glycolytic enzymes (Hk2, Pfkfb3 and Pkm) were also present in lower amounts in KO cells. Often, changes in ECAR are compensated by increased OCR but this was not the case in LRRC8A KO cells. Energy availability is the primary regulator of OCR. Despite a reduced ΔΨm, ADP/ATP and mitochondrial mass were unaltered. Thus, lower OCR is unlikely to reflect mitochondrial dysfunction. We hypothesize that low energy production in KO cells is related to reduced energy requirements for reduction of oxidized proteins and lipids, and for support of inflammation, proliferation and migration.

Lipolysis was the second most significantly altered pathway identified by KEGG analysis (Fig. 2A). Lipases are activated by cAMP to release free fatty acids (FFAs) from stored triglycerides to provide substrate for beta-oxidation. AKT activation by insulin blocks this process by phosphorylating and activating phosphodiesterase 3B (Pde3b, 11.3X increased, p < 0.01) which degrades cAMP. Expression of genes that promote cAMP production such as Ptgs2 (COX2) and Adcy8 (Adenylate cyclase 8) were reduced, but so was Ptger3, the prostaglandin E2 receptor which inhibits adenylate cyclase. Because FFAs are potent pro-inflammatory agents and promote atherosclerosis^64^ additional investigation into the status of lipolysis and fatty acid oxidation in LRRC8A KO cells is warranted.

AKT activation by insulin is impaired in LRRC8A KO adipocytes^65^ and skeletal muscle cells^45^, and PIP3 kinase and AKT activation by angiotensin II is disrupted in KO VSMCs^66^. PPI analysis identified EGFR as the top protein association hub. The insulin receptor and EGFR tyrosine kinases share key associated proteins (e.g., Irs1, Grb2), signaling pathways (PI3K/Akt and Ras/Erk)^67^, and Nox-dependance^68^. LRRC8A also associates with Grb2^69^, Nox1^17^, and Nox2^70^ and oxidants from Nox1 promote phosphorylation and transactivation of the EGFR^71^. EGFR activation (pY1068) was reduced in KO cells both at rest and in response to EGF. Activated EGFR associates with IRS1 which turns on PIP3 kinase, producing PIP3 which drives membrane association of AKT, mTORC2 and PDK1. This promotes AKT activation by phosphorylation at S473 and T308. The observed reduction in phosphorylation at these sites is therefore consistent with reduced PIP3 production due to impaired EGRF activation in the setting of reduced ROS.

### Inflammation and Senescence

The RNAseq data supported our prior observation of suppressed NF-κB activation by TNFα in cells treated with siRNA targeting VRACs^16,17^, as did luciferase reporter assays in KO cells (Fig. 5A). Reduced NF-κB and Hif-1α activity is consistent with less inflammation, and delayed senescence - a state characterized by growth arrest and the senescence-associated secretory phenotype (SASP)^72^. It is associated with aging-induced telomere shortening^31^ (replicative senescence), with oxidant stress^32,72,73^ (premature senescence), and with hypertension, atherosclerosis, and cardiovascular risk^74^. SASP genes were consistently under-expressed in KO VSMCs (Suppl. Table 1). β-gal, a classic marker of senescence, had reduced mRNA (Suppl. Table 1), protein expression (Fig. 5B), and activity (Fig. 5C) in KO cells. Reduced inflammation may explain longer telomeres of KO VSMCs. Consistent with these *in vitro* findings, VSMC-specific LRRC8A KO mice exposed to AngII infusion were protected from senescence as reflected by aortic β-gal^75^. We saw this again in the setting of atherosclerosis (Fig. 6B).

Surprisingly, selective KO of LRRC8A in endothelial cells was associated with accelerated senescence and vascular aging^76^. How can these opposite effects be reconciled? As discussed, Nox1-derived O_2_**^·^**^-^ drives VSMC inflammatory signaling. It can also lead to uncoupling of endothelial eNOS^77^. However, in endothelial cells laminar shear activates Nox2, which also interacts with LRRC8A^70^. These ROS promote eNOS phosphorylation and nitric oxide production^77^, and enhance cell survival^78^. Thus, the two cell types may interpret LRRC8A-dependent redox signals very differently.

### Atherosclerosis

The contribution of VSMCs to atheroma formation requires VSMC phenotypic switching from contractile to secretory. This is followed by migration to the intima and transformation into LDL-accumulating foam cells^28^. Disease progression is Nox1-dependent^79^ and characterized by oxidative stress, and increased glycolysis, proliferation, migration, and senescence^74^. Both KO, and importantly LRRC8A HET mice, were resistant to hypercholesterolemia-induced aortic atheroma formation. Expression of LRRC8A was correlated with markers of inflammation including ICAM and VCAM.

OxLDL promotes atherogenesis by activating LOX-1 and CD36 scavenger receptors^47,80^ that trigger ROS production by both Nox1 and mitochondria, and activate NF-κB. LOX-1 and CD36 abundance were not altered in aortae from KO mice, but uptake of OxLDL was significantly reduced in cultured KO VSMCs. Thus, resistance to atherosclerosis in KO mice may be related to impaired function of these receptors, as is the case with other Nox1-dependent inflammatory signals that drive the process. A connection to LRRC8A was supported by the finding that three structurally unrelated VRAC inhibitors reduced OxLDL uptake in WT VSMC to a degree like that seen in KO cells. This demonstrates that LRRC8A function, and potentially the ROS produced in response to LOX-1 or CD36 activation, is required for receptor endocytosis and OxLDL uptake. Importantly, LRRC8A KO did not alter uptake of non-oxidized LDL by the LDLR, which does not activate Nox1.

We previously assessed vascular reactivity in mesenteric arteries from VSMC-specific LRRC8A KO mice exposed to TNFα *in vitro* or to AngII hypertension *in vivo*^21,35^. In freshly isolated KO vessels, contractile responses to KCl and PE were unaltered while relaxation to both ACh and SNP were augmented, while in both models of inflammatory disease the vessels were protected from vascular dysfunction. In another pro-inflammatory setting of hypercholesterolemia, contractility to PE and NE were increased in WT but not in KO or HET vessels. Protected dilation to ACh was again seen in female but not in male mice. This sex difference is in accordance with a prior demonstration of impaired responses to ACh only in female ApoE^-/-^ mice^81,82^. We attributed vasomotor changes in VSMC-specific LRRC8A KO mice to reduced RhoA activity and hypothesized that this was due to the loss of activating effects of O2**^·^**^-^ on the small GTPase positioned in proximity to oxidants in a multi-protein complex organized by MPRIP. Here we assessed expression of key proteins in this complex and found reduced expression of MYPT1 and CPI17, both of which are capable of impairing activity of MLCP and increasing calcium sensitivity.

### Summary

Impaired AKT activation by insulin has been well documented in LRRC8A KO cells^65^. We now report similar defects in EGFR signaling, suggesting a shared defect in protein tyrosine kinase signaling, that may be attributable to reduced oxidant influx. The integrated downstream effects of these changes include lower Hif-1α and NF-κB activity, reduced inflammation, proliferation and migration all of which lower energy demand, reducing both ECAR and OCR while maintaining energy availability.

These signaling defects protect VSMCs in the pro-inflammatory environment of hypercholesterolemia. Agonist-triggered cytoplasmic oxidant signals may be required for endocytosis of the LOX-1 and CD36 scavenger receptors as they are for TNFR1^17^. LRRC8A-dependance of OxLDL uptake is a key novel finding that will require detailed investigation to determine how VRACs and ROS contribute to this process. Collectively, these data define LRRC8A as an exciting potential target for selective inhibition of ROS-dependent pathophysiology.

## ACKNOWLEDGMENTS

We thank Dr. Benjamin Crawford for his expert assistance with flow cytometry data acquisition.

## FOOTNOTE

## Nonstandard Abbreviations and Acronyms

Ach: Acetylcholine
ApoE: Apolipoprotein E
ApoE⁻/⁻: Apolipoprotein E–knockout
ASK1: Apoptosis signal–regulating kinase 1
β-Gal: Β-galactosidase
CBX: Carbenoxolone
CPI-17: PKC-potentiated inhibitor protein of PP1 (17 kda)
DHE: Dihydroethidium
ECAR: Extracellular acidification rate
EGFR: Epidermal growth factor receptor
FACS: Fluorescence-activated cell sorting
HCT116: Human colorectal carcinoma cell line HCT-116
HET: Heterozygous
HIF-1α: Hypoxia-inducible factor-1α
HK2: Hexokinase 2
HO-1: Heme oxygenase-1
HRE: Hypoxia response element
ICAM: Intercellular adhesion molecule-1
IGF1R: Insulin-like growth factor-1 receptor
IRS1: Insulin receptor substrate-1
KO: Knockout
LDL: Low-density lipoprotein
LDLR: Low-density lipoprotein receptor
LOX-1: Lectin-like oxidized LDL receptor-1
LRRC8A/C/D: Leucine-rich repeat–containing 8A/8C/8D
MDA: Malondialdehyde
MLC: Myosin light chain
MLCP: Myosin light chain phosphatase
MYPT1: Myosin phosphatase target subunit-1
NE: Norepinephrine
Nox1: NADPH oxidase 1
Nrf2: Nuclear factor erythroid 2–related factor 2
OCR: Oxygen consumption rate
OxLDL: Oxidized low-density lipoprotein
PCNA: Proliferating cell nuclear antigen
Pdk1: 3-phosphoinositide-dependent protein kinase-1
Pfkfb3: 6-phosphofructo-2-kinase/fructose-2
PIP₃: Phosphatidylinositol 3,4,5-trisphosphate
PKM: Pyruvate kinase M
PPI: Protein–protein interaction
Rap1: Ras-associated protein 1
RNAseq: RNA sequencing
ROS: Reactive oxygen species
ROSstar550: Cell-permeant fluorescent ROS probe
Rps4l: Ribosomal protein S4-like
ROCK: Rho-associated coiled-coil containing protein kinase
SA-β-gal: Senescence-associated β-galactosidase
SASP: Senescence-associated secretory phenotype
SNP: Sodium nitroprusside
SOD3: Extracellular superoxide dismutase
TAM: Tamoxifen
TMRE: Tetramethylrhodamine ethyl ester
TNFα: Tumor necrosis factor-α
VDAC1: Voltage-dependent anion channel 1
VCAM: Vascular cell adhesion molecule-1
VRAC: Volume-regulated anion channel
VSMC: Vascular smooth muscle cell
WT: Wild type
ΔΨm: Mitochondrial membrane potential

