## Supplemental Materials for "Smooth muscle LRRC8A knockout reduces O_2_^·-^ influx, inflammation, senescence and atherosclerosis"

### Supplementary File

### Supplemental Figure 1

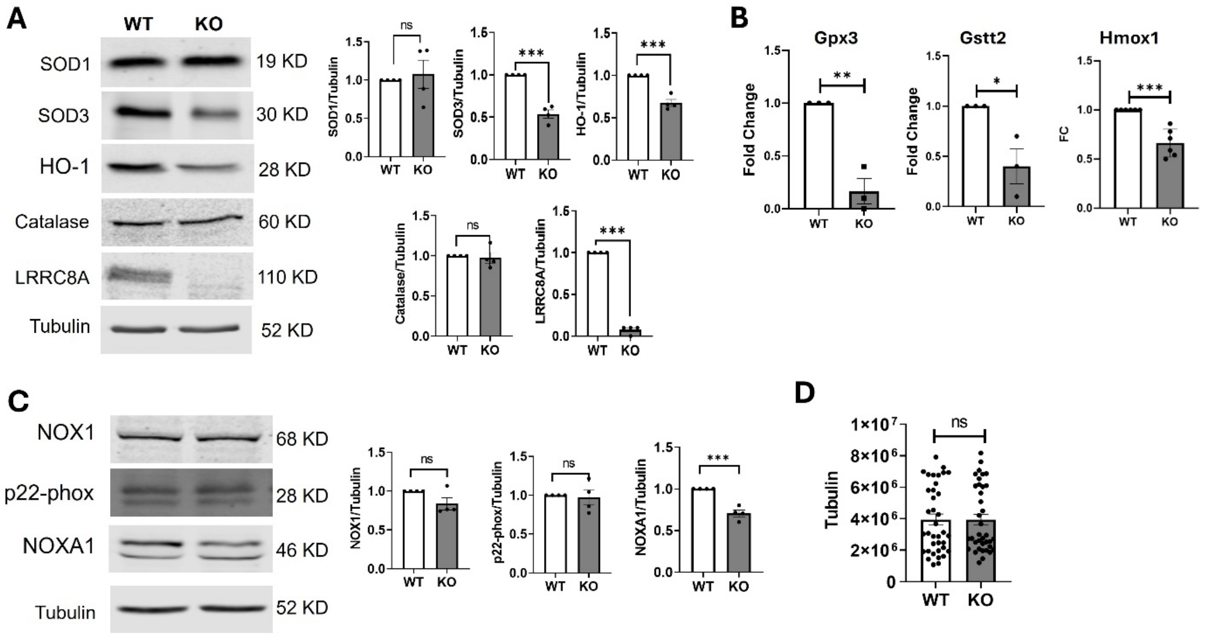

**Supplemental Figure 1.** Antioxidant enzymes and Nox1 complex components in primary aortic cultured VSMCs from WT and LRRC8A KO mice. **A,** Immunoblotting for SOD1, SOD3, HO-1, catalase, LRRC8A, and α-tubulin normalized to α-tubulin. **B,** qPCR for Gpx3, Gstt2, and Hmox1 in WT and KO VSMCs normalized to housekeeping transcripts and expressed relative to WT. **C,** Immunoblotting for NOX1, p22-phox, and NOXA1 normalized to α-tubulin. **D,** Loading-control verification; total α-tubulin signal per sample across WT and KO preparations of cultured cells. Data represents SEM, dots represent individual replicates. *$P$<0.05, **$P$<0.01, ***$P$<0.001.

### Supplemental Figure 2

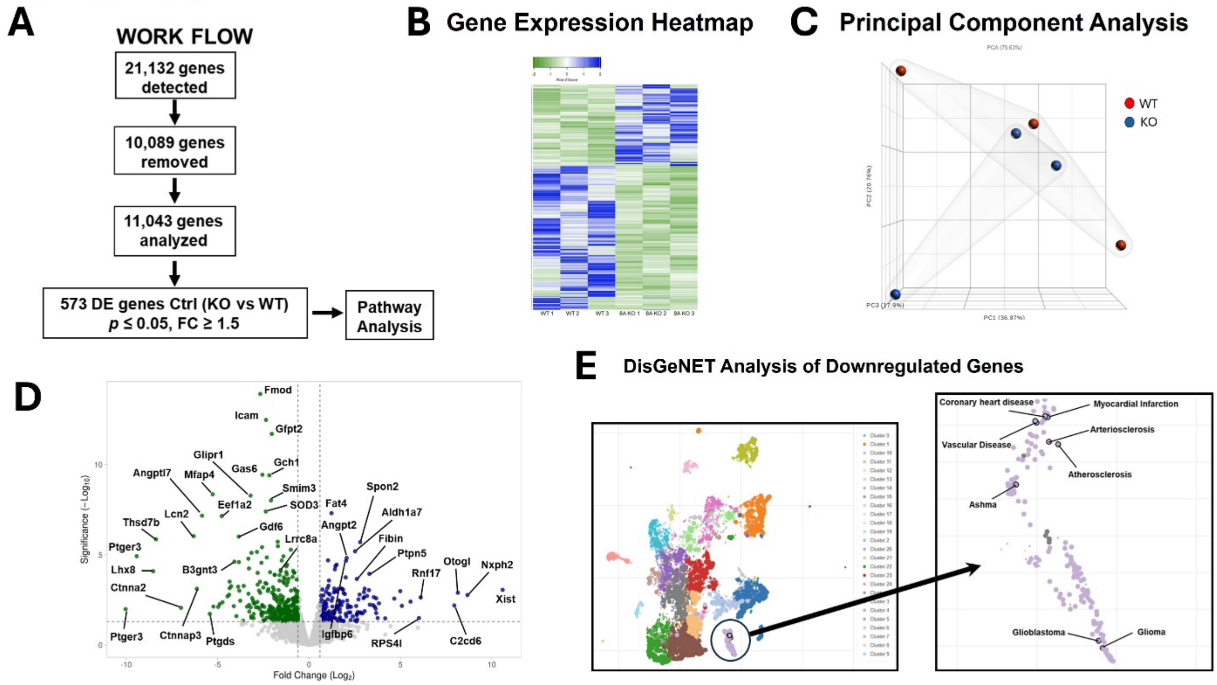

**Supplemental Figure 2. RNA-seq workflow and global differential expression in LRRC8A-deficient VSMCs.**
**A,** Workflow for bulk RNA-seq from primary cultured aortic VSMCs (WT and LRRC8A KO; $n = 3$ replicates of cells pooled from 3 animals per genotype). After filtering low-abundance features, 11,043 genes were retained; 573 differentially expressed (DE) genes (KO vs WT) met preset thresholds ($P \leq 0.05$; |fold change|≥ 1.5) and were utilized for pathway analysis. **B,** Hierarchical clustering of the DE gene set shows clear segregation by genotype, with coherent down- and up-regulated blocks across all biological replicates. **C,** Principal component analysis (PCA) of all retained genes showing sample separation by genotype. **D,** Genome-wide differential expression (KO vs WT) displayed as $\log_2$ fold-change versus $-\log_{10}(p)$, representative genes are annotated. **E,** DisGeNET/Enrichr disease-association analysis of downregulated genes: cluster map (left) and zoomed view of representative enriched disease terms (right). Details of library preparation, alignment/quantification, normalization, and enrichment tools/parameters are provided in Methods. Abbreviations: WT, wild type; KO, LRRC8A knockout; DE, differentially expressed.

### Supplemental Figure 3

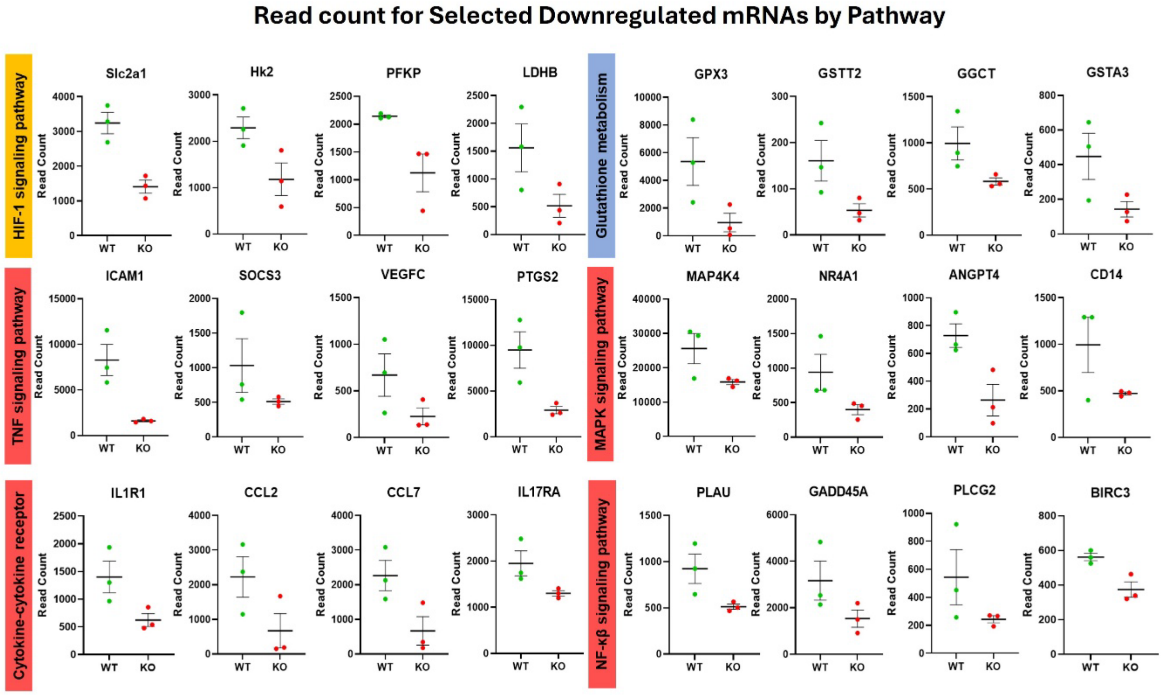

**Supplementary Figure 3. RNA-seq read counts for selected transcripts grouped by KEGG pathways.** Normalized RNA-seq read counts from primary aortic VSMCs (WT and LRRC8A-KO; n = 3 per genotype) are shown as scatter plots with mean ± SEM error bars. *P*-value of all those genes are below 0.05. Genes are organized by pathway: HIF-1 signaling (Slc2a1, Hk2, Pfkp, Ldhb), Glutathione metabolism (Gpx3, Gstt2, Ggct, Gsta3), TNFα signaling (Icam1, Socs3, Vegfc, Ptgs2) MAPK signaling (Map4k4, Nr4a1, Angpt4, Cd14), Cytokine-cytokine receptor interaction (Il1r1, Ccl2, Ccl7, Il17ra), and NF-κB signaling (Plau, Gadd45a, Plcg2, Birc3). Read-count normalization and differential-expression criteria are described in Methods. Abbreviations: WT, wild type; KO, LRRC8A knockout

### Supplemental Figure 4

#### Pathways in Upregulated Genes (Kegg)

| Sr. No. | Term | P-value | Combined Score | Genes |
|---|---|---|---|---|
| 1 | Rap1 signaling pathway | 0.00 | 28.05 | GNAO1;EFNA3;SIPA1L1;ANGPT2;HGF;EVL;THBS1;PFN2 |
| 2 | Terpenoid backbone biosynthesis | 0.00 | 108.27 | FDPS;MVD;ACAT2 |
| 3 | Gap junction | 0.00 | 39.51 | GJA1;GUCY1A1;TUBB2B;GUCY1B1;TUBA1A |
| 4 | Purine metabolism | 0.01 | 19.62 | GUCY1A1;GUCY1B1;PDE3B;AK5;PDE8B |
| 5 | Thiamine metabolism | 0.01 | 73.54 | AK5;ACP1 |
| 6 | Morphine addiction | 0.01 | 20.75 | GNAO1;GRK3;PDE3B;PDE8B |
| 7 | Circadian entrainment | 0.02 | 18.46 | GNAO1;GUCY1A1;GUCY1B1;GRIA4 |
| 8 | Proteoglycans in cancer | 0.02 | 12.90 | COL1A1;FZD2;HGF;PDCD4;ESR1;THBS1 |
| 9 | Regulation of lipolysis in adipocytes | 0.02 | 24.15 | IRS1;PDE3B;NPY1R |
| 10 | Cushing syndrome | 0.02 | 13.70 | FZD2;CDKN2A;CYP11B1;AHR;PDE8B |
| 11 | Protein digestion and absorption | 0.02 | 16.51 | COL1A1;COL3A1;COL24A1;COL5A1 |
| 12 | Long-term depression | 0.02 | 20.78 | GNAO1;GUCY1A1;GUCY1B1 |
| 13 | Wnt signaling pathway | 0.02 | 11.92 | FZD2;DAAM2;LGR6;DKK2;LGR4 |
| 14 | Renin secretion | 0.03 | 16.26 | GUCY1A1;GUCY1B1;PDE3B |
| 15 | Axon guidance | 0.03 | 9.84 | NRP1;EFNA3;FES;EPHB3;RGMA |
| 16 | Non-small cell lung cancer | 0.03 | 15.07 | CDKN2A;HGF;RARB |
| 17 | Platelet activation | 0.03 | 11.57 | COL1A1;GUCY1A1;COL3A1;GUCY1B1 |
| 18 | Glyoxylate and dicarboxylate metabolism | 0.03 | 24.47 | ACSS2;ACAT2 |
| 19 | Propanoate metabolism | 0.04 | 19.95 | ACSS2;ACAT2 |
| 20 | Insulin signaling pathway | 0.05 | 9.49 | SOCS2;PPP1R3C;IRS1;PDE3B |

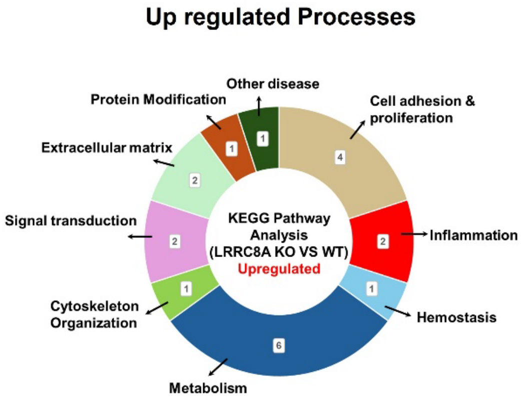

**Supplementary Figure 4. KEGG pathway analysis of upregulated genes in LRRC8A-KO VSMCs. A,** Table of the top KEGG pathways enriched among transcripts increased in KO versus WT analysis (bulk RNAseq; n = 3 per genotype). Columns list the pathway term, nominal *P*, combined score and representative member genes. **B,** Category wheel summarizing broader biological processes represented by the upregulated pathways. Gene selection thresholds and enrichment parameters are described in Methods. Abbreviations: WT, wild type; KO, LRRC8A knockout

### Supplemental Figure 5

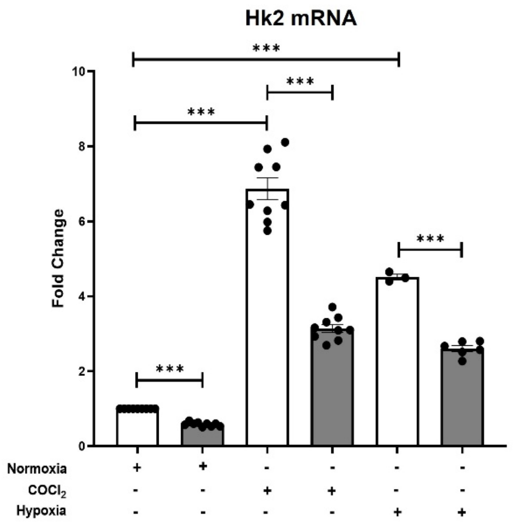

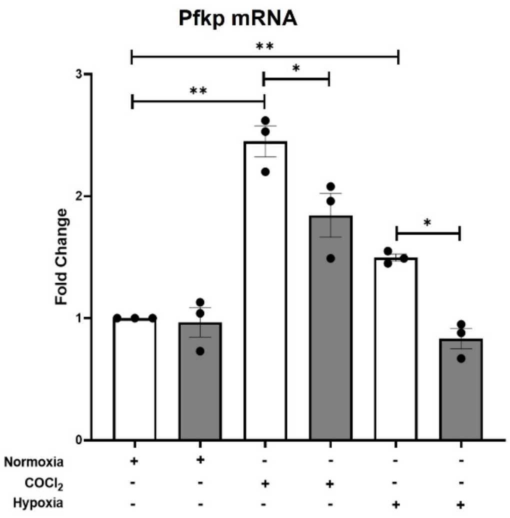

**Supplementary Figure 5. HIF-1α-responsive glycolytic transcripts under hypoxic stimuli.** qPCR for Hk2 and Pfkp mRNA in primary aortic VSMCs (WT and LRRC8A KO) cultured under normoxia, $CoCl_2$ (300 µM, 6 h), or hypoxia (1% $O_2$, 24 h). Values are normalized to housekeeping GAPDH gene and expressed relative to WT in normoxia. Data are expressed as mean ± SEM with individual biological replicates shown. For HK2: normoxia comparisons were evaluated by one-sample *t*-test against the theoretical mean of 1; $CoCl_2$ and hypoxia WT vs KO comparisons used unpaired *t*-tests (n = 9 for $CoCl_2$; n = 6 for hypoxia). For PFKP: n = 3 per condition, and tests were performed as for HK2.

### Supplemental Figure 6

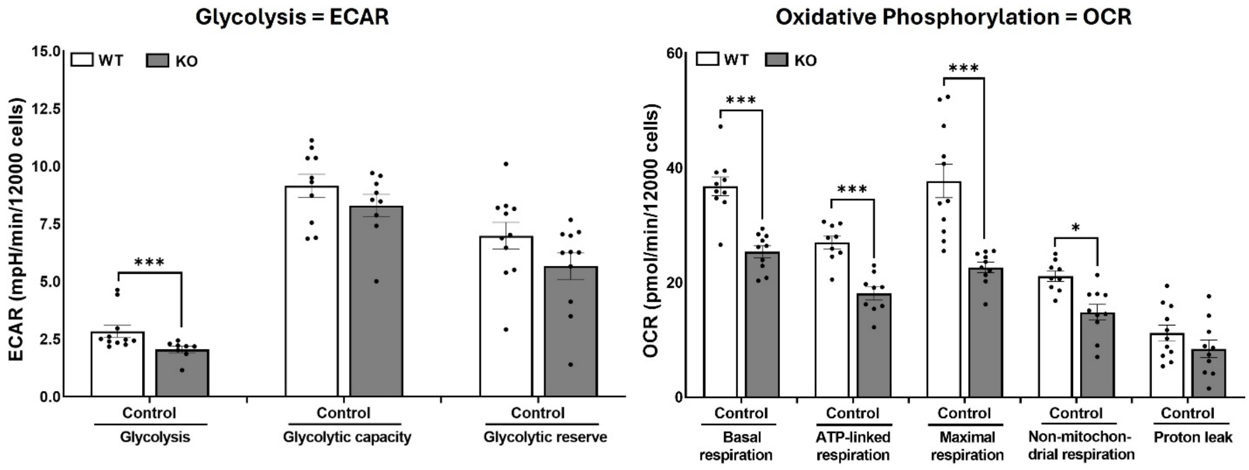

**Supplementary Figure 6. Summary of extracellular flux parameters in WT and LRRC8A-KO VSMCs.** *Left*, ECAR parameters from the glycolysis stress test (glucose → oligomycin → 2-deoxy-D-glucose): basal glycolysis, glycolytic capacity, and glycolytic reserve. Right, OCR parameters from the mitochondrial stress test (oligomycin → FCCP → antimycin A/rotenone): basal respiration, ATP-linked respiration, maximal respiration, non-mitochondrial respiration, and proton leak. Points represent independent biological preparations; bars are mean ± SEM (n = 10 per genotype). Analyzed using one-way ANOVA with Tukey's post-hoc test). Abbreviations: WT, wild type; KO, LRRC8A knockout; ECAR, extracellular acidification rate; OCR, oxygen consumption rate. $*P<0.05$, $**P<0.01$, $***P<0.001$.

### Supplementary Figure 7

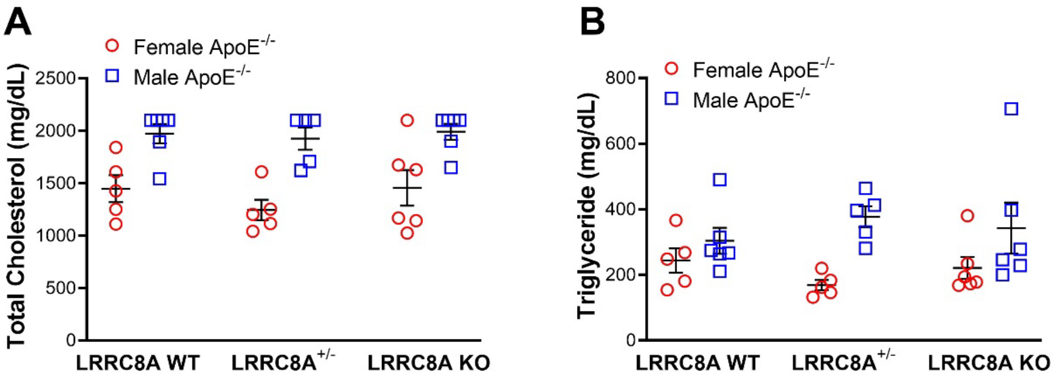

**Supplementary Figure 7. Serum lipids in ApoE$^{-/-}$ mice were not affected by VSMC-specific LRRC8A KO.** Plasma lipids in female (red circles) and male (blue squares) ApoE$^{-/-}$ mice with LRRC8A WT, HET, or KO genotypes. **A,** Total cholesterol (mg/dL). Values above 2000 were offscale and are demoted as 2000. **B,** Triglycerides (mg/dL). Symbols denote individual mice, graphs show mean ± SEM. Analyses by one-way ANOVA showed no changes (male n = 5-6, female n = 5-6).

### Supplemental Table 1

| Symbol | Name | Fold | P |
|---|---|---|---|
| **Contractile** | | | |
| Acta2 | SM actin | -1.55 | 0.6124 |
| Actn1 | Alpha actinin | 1.05 | 0.7892 |
| Actg2 | Smooth muscle gamma actin | 1.03 | 0.9778 |
| Aebp1 | Adipocyte enhancer binding protein 1 | -1.57 | 0.1383 |
| Cnn2 | Calponin 2 | 1.01 | 0.9797 |
| Cald1 | Caldesmon | -1.53 | 0.1791 |
| Csrp1 | Cysteine and glycine rich protein 1 | -1.47 | 0.3587 |
| Dmd | Dystrophin | -1.21 | 0.6069 |
| Glmn | Glomulin | 1.24 | 0.3793 |
| Hdac8 | Histone deacetylase 8 | -1.04 | 0.8620 |
| Itga8 | Integrin alpha 8 | -1.36 | 0.7068 |
| Itga1 | Integrin alpha 1 | -2.29 | 0.0040 |
| Myh11 | Myosin Heavy Chain | -11.88 | 0.0005 |
| Notch3 | Notch receptor 3 | -1.26 | 0.6075 |
| Ptk2 | Protein Tyrosine Kinase 2 | 1.15 | 0.5934 |
| Smtn | Smoothelin | -1.19 | 0.2399 |
| Srf | Serum response factor | -1.10 | 0.6718 |
| Tagln | Smooth muscle 22alpha | -1.17 | 0.8596 |
| Tpm1 | Alpha tropomysin | -1.04 | 0.8980 |
| Tpm2 | Beta tropomysin | -1.66 | 0.2305 |
| Vcl | Vinculin | -1.51 | 0.1961 |
| **Lmod1 | Leiomodin 1 | -4.00 | 0.0448 |
| **Cnn1 | Calponin 1 | -2.97 | 0.3666 |
| **Synthetic** | | | |
| Ereg | Epiregulin | -2.30 | 0.0141 |
| Rbp1 | Retinol binding protein | 1.60 | 0.2822 |
| Vim | Vimentin | 1.15 | 0.4957 |
| **col1a1 | Collagen Type I Alpha 1 Chain | 1.52 | 0.0212 |
| **Myh10 | Myosin Heavy Chain 10 | 1.31 | 0.2503 |
| **Spp1 | Secreted Phosphoprotein 1 | -2.02 | 0.1512 |
| **Klf4 | KLF Transcription Factor 4 | 1.05 | 0.8761 |
| **Lgals3 | Galectin 3 | 1.65 | 0.2438 |
| **Fibroblast** | | | |
| Lum | Lumican | -3.87 | 0.3685 |
| Bgn | Biglycan | -1.10 | 0.6057 |
| Dcn | Decorin | 1.57 | 0.3974 |
| **Adipose** | | | |
| Dio2 | Iodothyronine deiodinase 2 | -1.81 | 0.4664 |
| Ppargc1 | peroxisome proliferative activated receptor, gamma, coactivator 1 alpha | ND | |
| Ucp1 | Uncoupling protein 1 | ND | |
| **Osteogenic** | | | |
| Alpl | Alkaline phosphatase liver/bone | -8.64 | 0.0025 |
| Runx2 | Runt-related transcription factor-2 | -1.09 | 0.6631 |
| Spp1 | Osteopontin | -2.02 | 0.1512 |
| Tnfrsf11b | TNF receptor superfamily, member 11b (osteoprotegerin) | -1.16 | 0.7408 |
| **Mesenchymal** | | | |
| Cd34 | Cluster of Differentiation 34 | -2.52 | 0.3650 |
| Cd44 | Cluster of Differentiation 44 | -1.38 | 0.1031 |
| Ly6a | Lymphocyte antigen 6 family member a | 1.68 | 0.1494 |
| **Macrophage** | | | |
| Adgre1 | adhesion G protein-coupled receptor E1 | ND | |
| Itgam | Integrin alpha M | ND | |
| Cd68 | Macrosialin | -1.70 | 0.3353 |
| Lgals3 | lectin, galactose binding, soluble 3 | 1.65 | 0.2438 |
| Lamp2 | Lysosomal-associated membrane protein-2 | -1.06 | 0.8251 |
| Vcam1 | Vascular cell adhesion molecule 1 | -1.04 | 0.9344 |

| Symbol | Name | Fold | P |
|---|---|---|---|
| **Senescence Markers** | | | |
| *Cytokines and Chemokines* | | | |
| Ccl2 | CC motif chemokine ligand 2 (MCP-1) | -3.32 | 0.0380 |
| Ccl7 | CC motif chemokine ligand 7 (MCP-3) | -3.38 | 0.0120 |
| Ccl8 | MCP-2, monocyte chemoattractant protein 2 | -1.62 | 0.5200 |
| Ccl9 | CC motif chemokine ligand 9 (Mip1gamma) | -1.15 | 0.8500 |
| Cxcl1 | C-X-C Motif Chemokine Ligand 1 | -1.79 | 0.2700 |
| Cxcl12 | C-X-C Motif Chemokine Ligand 12 (SDF-1) | 1 | 0.9900 |
| Il6 | Interleukin 6 | -5.94 | 0.0153 |
| Il15 | Interleukin 15 | 1.97 | 0.6800 |
| Mif | Macrophage migration inhibitory factor | 1 | 0.9900 |
| *Growth Factors and Regulators* | | | |
| Cdkn2a | Cyclin dependent kinase inhibitor 2A (p16) | 1.83 | 0.0047 |
| Cdkn1a | Cyclin dependent kinase inhibitor 1A (p21) | -1.09 | 0.7066 |
| Stat3 | Signal transducer and activator of transcription 3 | -1.51 | 0.0644 |
| Vegfc | Vascular endothelial growth factor C | -2.96 | 0.0180 |
| Plgf | Placental growth factor | 1.02 | 0.9300 |
| Ngf | Nerve growth factor | -1.91 | 0.0600 |
| Igfbp2 | Insulin-like growth factor binding protein 2 | -20.66 | 0.0016 |
| Igfbp3 | Insulin-like growth factor binding protein 3 | -14.04 | 0.0010 |
| Igfbp6 | Insulin-like growth factor binding protein 6 | 4.04 | 0.0000 |
| Ereg | Epiregulin | -2.3 | 0.0140 |
| Nrgl | Neuregulin | 1.25 | 0.5300 |
| Fgf2 | Fibroblast growth factor 2 (beta FGF) | -2.32 | 0.4700 |
| Hgf | Hepatocyte growth factor | 3.38 | 0.0460 |
| *Proteases and Regulors* | | | |
| Mmp13 | Matrix metalloproteinase 13 | -5.33 | 0.0970 |
| Mmp16 | Matrix metalloproteinase 16 | -3.82 | 0.0200 |
| Mmp27 | Matrix metalloproteinase 27 | -2.78 | 0.0980 |
| Mmp | Mmp2, 3, 11, 14, 15, 19, 23, 28 | | NS |
| Timp1 | | 1.02 | 0.9500 |
| Timp2 | | 1.03 | 0.9300 |
| Serpine1 | Plasminogen activator inhibitor-1 (PAI-1) | -1.80 | 0.0342 |
| Ctsb | Cathepsin B | -1.22 | 0.4500 |
| *Shed receptors or ligands* | | | |
| Tlr4 | Toll-like receptor 4 | -1.31 | 0.5278 |
| Icam1 | Intercellular adhesion molecule 1 | -5.10 | 0.0000 |
| Tnfr1a | TNF receptor type I | -1.23 | 0.1900 |
| Tnfr2 | TNF receptor type II | -1.48 | 0.2900 |
| Fas | Fas cell surface death receptor (Tnfrsf6) | -1.32 | 0.1400 |
| Egfr | Epidermal growth factor receptor | -1.49 | 0.1200 |
| Plaur | Urokinase | -1.65 | 0.1500 |
| Il6st | Interleukin 6 cytokine family signal transducer | -1.32 | 0.3400 |
| *Other* | | | |
| Glb1 | Lysosomal β-galactosidase 1 | -1.72 | 0.0008 |

| Symbol | Name | Fold |
|---|---|---|
| **Autophagy** | | |
| Akt1 | Rac(Rho family)-alpha serine/threonine-protei | 1.23 |
| Akt2 | Rac-beta serine/threonine-protein kinase | 1.24 |
| Ambra1 | Activating molecule in Beclin1-regulated autop | 1.01 |
| Atg3 | Autophagy related 3 | -1.05 |
| Atg4b | Cysteine protease Atg4b | -1.18 |
| Atg5 | Autophagy related 5 | -1.16 |
| Atg7 | Autophagy related 7 | -1.32 |
| Atg9a | Autophagy related 9A | -1.02 |
| Atg12 | Autophagy related 12 | -1.14 |
| Atg13 | Autophagy related 13 | -1.09 |
| Atg14 | Autophagy related 14 | -1.08 |
| Atg16l1 | Autophagy related 16 like 1 | -1.34 |
| Atg101 | Autophagy related 101 | -1.09 |
| Becn1 | Beclin-1 | -1.19 |
| Bnip3 | Bcl2/adenovirus E1B 19 kDa protein-interacting | -1.05 |
| Bnip3l | Bcl2/adenovirus E1B 19 kDa protein-interacting | -1.13 |
| Ctsd | Cathepsin D | -1.24 |
| Gabarap | GABA type A receptor-associated protein | -1.09 |
| Hif1a | Hyoxia-inducible factor alpha | -1.36 |
| Hspa8 | Heat shock 70 kDa protein 8 | 1.08 |
| Lamp2 | Lysosomal-associated membrane protein-2 | -1.06 |
| Map1lc3a | Microtubule-associated proteins 1A/1B light ch | 1.20 |
| Map1lc3b | Microtubule-associated proteins 1A/1B light ch | 1.03 |
| Mtor | Mammalian target of rapamycin | -1.27 |
| Nbr1 | Neighbor of BRCA1 gene 1 protein | -1.33 |
| Nfe2l2 | Nuclear factor erythroid 2-related factor 2 (Nrf2 | -1.11 |
| Optn | Optineurin | -1.38 |
| Pik3c3 | Phosphatidylinositol 3-kinase catalytic subunit | -1.08 |
| Pik3r4 | Phosphatidylinositol 3-kinase catalytic subunit | -1.18 |
| Pink1 | Pten-induced kinase 1 | -1.26 |
| Prkaa1 | 5'-AMP-activated protein kinase catalytic subun | -1.08 |
| Rb1cc1 | RB1-inducible coiled-coil protein 1 | 1.08 |
| Sqstm1 | Sequestosome-1 | -1.41 |
| Stx17 | Syntaxin 17 | 1.01 |
| Tax1bp1 | Tax1-binding protein 1 | -1.26 |
| Tex264 | Testis expressed 264 ER-phagy receptor | -1.32 |
| Ulk1 | Serine/threonine-protein kinase Ulk1 | 1.11 |
| Wipi1 | WD repeat domain phosphoinositide-interactin | 1.04 |
| Zfyve1 | Zinc finger FYVE domain-containing protein 1 | 1.15 |
| **Apoptosis** | | |
| Casp2 | Caspase 2 | -1.15 |
| Casp3 | Caspase 3 | 1.23 |
| Casp7 | Caspase 7 | 1.27 |
| Casp8 | Caspase 8 | 1.09 |
| Casp9 | Caspase 9 | -1.15 |
| Cycs | Cytochrome C | -1.01 |
| Fas | Fas cell surface death receptor | -1.32 |
| Bax | Bcl-2 associated X | 1.18 |
| Bak1 | Bcl-2 homologous antagonist/killer | 1.21 |
| Bcl2 | B -cell lymphoma 2 | 1.21 |
| Trp53 | Transformation related protein 53 (p53) | -1.13 |

**Table 1. Expression of genes associated with senescence are under-expressed in LRRC8A KO VSMCs.** Gene name, fold change and P-value are provided for markers of contractile vs. secretory phenotype, or altered cell type (fibroblast, adipocyte, osteocyte, mesenchymal, macrophage) in column 1. Markers or senescence are listed in column 2. Markers of autophagy or apoptosis are provided in column 3. Genes with significantly reduced expression are highlighted in green. Those with increased expression are highlighted in red.

### SUPPLEMENTARY FIGURE LEGENDS

**Supplemental Figure 1.** Antioxidant enzymes and Nox1 complex components in primary aortic cultured VSMCs from WT and LRRC8A KO mice. **A,** Immunoblotting for SOD1, SOD3, HO-1, catalase, LRRC8A, and α-tubulin normalized to α-tubulin. **B,** qPCR for Gpx3, Gstt2, and Hmox1 in WT and KO VSMCs normalized to housekeeping transcripts and expressed relative to WT. **C,** Immunoblotting for NOX1, p22-phox, and NOXA1 normalized to α-tubulin. **D,** Loading-control verification; total α-tubulin signal per sample across WT and KO preparations of cultured cells. Data represents SEM, dots represent individual replicates. *$P<0.05$, **$P<0.01$, ***$P<0.001$.

**Supplemental Figure 2. RNA-seq workflow and global differential expression in LRRC8A-deficient VSMCs.**

**A,** Workflow for bulk RNA-seq from primary cultured aortic VSMCs (WT and LRRC8A KO; *n* = 3 replicates of cells pooled from 3 animals per genotype). After filtering low-abundance features, 11,043 genes were retained; 573 differentially expressed (DE) genes (KO vs WT) met preset thresholds ($P \leq 0.05$; |fold change|≥ 1.5) and were utilized for pathway analysis. **B,** Hierarchical clustering of the DE gene set shows clear segregation by genotype, with coherent down- and up-regulated blocks across all biological replicates. **C,** Principal component analysis (PCA) of all retained genes showing sample separation by genotype. **D,** Genome-wide differential expression (KO vs WT) displayed as $\log_2$ fold-change versus $-\log_{10}(p)$, representative genes are annotated. **E,** DisGeNET/Enrichr disease-association analysis of downregulated genes: cluster map (left) and zoomed view of representative enriched disease terms (right). Details of library preparation, alignment/quantification, normalization, and enrichment tools/parameters are provided in Methods. Abbreviations: WT, wild type; KO, LRRC8A knockout; DE, differentially expressed.

**Supplementary Figure 3. RNA-seq read counts for selected transcripts grouped by KEGG pathways.** Normalized RNA-seq read counts from primary aortic VSMCs (WT and LRRC8A-KO; n = 3 per genotype) are shown as scatter plots with mean ± SEM error bars. *P*-value of all those genes are below 0.05. Genes are organized by pathway: HIF-1 signaling (Slc2a1, Hk2, Pfkp, Ldhb), Glutathione metabolism (Gpx3, Gstt2, Ggct, Gsta3), TNFα signaling (Icam1, Socs3, Vegfc, Ptgs2) MAPK signaling (Map4k4, Nr4a1, Angpt4, Cd14), Cytokine-cytokine receptor interaction (Il1r1, Ccl2, Ccl7, Il17ra), and NF-κB signaling (Plau, Gadd45a, Plcg2, Birc3). Read-count normalization and differential-expression criteria are described in Methods. Abbreviations: WT, wild type; KO, LRRC8A knockout

**Supplementary Figure 4. KEGG pathway analysis of upregulated genes in LRRC8A-KO VSMCs. A,** Table of the top KEGG pathways enriched among transcripts increased in KO versus WT analysis (bulk RNAseq; n = 3 per genotype). Columns list the pathway term, nominal *P*, combined score and representative member genes. **B,** Category wheel summarizing broader biological processes represented by the upregulated pathways. Gene selection thresholds and enrichment parameters are described in Methods. Abbreviations: WT, wild type; KO, LRRC8A knockout.

**Supplementary Figure 5. HIF-1α-responsive glycolytic transcripts under hypoxic stimuli.** qPCR for Hk2 and Pfkp mRNA in primary aortic VSMCs (WT and LRRC8A KO) cultured under normoxia, $CoCl_2$ (300 μM, 6 h), or hypoxia (1% $O_2$, 24 h). Values are normalized to housekeeping GAPDH gene and expressed relative to WT in normoxia. Data are expressed as mean ± SEM with individual biological replicates shown. For HK2: normoxia comparisons were evaluated by one-sample *t*-test against the theoretical mean of 1; $CoCl_2$ and hypoxia WT vs KO comparisons used unpaired *t*-tests (n = 9 for $CoCl_2$; n = 6 for hypoxia). For PFKP: n = 3 per condition, and tests were performed as for HK2.

**Supplementary Figure 6. Summary of extracellular flux parameters in WT and LRRC8A-KO VSMCs.** *Left*, ECAR parameters from the glycolysis stress test (glucose → oligomycin → 2-deoxy-D-glucose): basal glycolysis, glycolytic capacity, and glycolytic reserve. Right, OCR parameters from the mitochondrial stress test (oligomycin → FCCP → antimycin A/rotenone): basal respiration, ATP-linked respiration, maximal respiration, non-mitochondrial respiration, and proton leak. Points represent independent biological preparations; bars are mean ± SEM (n = 10 per genotype). Analyzed using one-way ANOVA with Tukey's post-hoc test). Abbreviations: WT, wild type; KO, LRRC8A knockout; ECAR, extracellular acidification rate; OCR, oxygen consumption rate. *$P$<0.05, **$P$<0.01, ***$P$<0.001.

**Supplementary Figure 7. Serum lipids in ApoE$^{-/-}$ mice were not affected by VSMC-specific LRRC8A KO.** Plasma lipids in female (red circles) and male (blue squares) ApoE$^{-/-}$ mice with LRRC8A WT, HET, or KO genotypes. **A,** Total cholesterol (mg/dL). Values above 2000 were offscale and are demoted as 2000. **B,** Triglycerides (mg/dL). Symbols denote individual mice, graphs show mean ± SEM. Analyses by one-way ANOVA showed no changes (male n = 5-6, female n = 5-6).

**Table 1. Expression of genes associated with senescence are under-expressed in LRRC8A KO VSMCs.** Gene name, fold change and P-value are provided for markers of contractile vs. secretory phenotype, or altered cell type (fibroblast, adipocyte, osteocyte, mesenchymal, macrophage) in column 1. Markers or senescence are listed in column 2. Markers of autophagy or apoptosis are provided in column 3. Genes with significantly reduced expression are highlighted in green. Those with increased expression are highlighted in red.

### SUPPLEMENTARY METHODS

#### Animal Studies and Ethics Statement

All animal procedures were conducted in strict accordance with the recommendations in the Guide for the Care and Use of Laboratory Animals of the National Institutes of Health and were approved by the Institutional Animal Care and Use Committee (IACUC) of Vanderbilt University Medical center (Protocol #M1600151). All studies were performed and reported in compliance with the ARRIVE guidelines. To generate mice for primary cell isolation, Lrrc8a$^{fl/fl}$ mice (C57B6/129 mixed background; a gift from Dr. Rajan Sah, Washington University) were crossed with transgenic mice expressing Cre recombinase under the control of the smooth muscle-specific α-actin promoter (SM22α-Cre; The Jackson Laboratory, Stock No: 017491). Wild Type (WT) and KO littermates underwent experimentation. Mice were housed in a specific pathogen-free, AAALAC-accredited facility on a 12-hour light/dark cycle with ad libitum access to water and standard chow. Genotyping was performed by PCR analysis of tail-tip DNA by TransnetYX, Inc. (Memphis, TN).

#### *Primary Vascular Smooth Muscle Cell (VSMC) Isolation and Culture*

Primary aortic VSMCs were isolated from thoracic aortae in 3 male WT or LRRC8A KO mice by the explant method. Briefly, thoracic aortae were excised and cleaned of adherent fat. The endothelial layer was removed by passing a pin through the lumen and the aortae were cut into 2 mm square sections in ice-cold PSS. The segments were placed in culture plates and maintained in Dulbecco's Modified Eagle Medium (DMEM) containing 30% fetal bovine serum (FBS) and 1% penicillin/streptomycin in a humidified incubator at 37 °C, 5% $CO_2$ atmosphere. After 1 week when cells had migrated out of the tissue fragments the explants were removed and the migrated cells maintained in DMEM supplemented with 10% FBS, 1% penicillin/streptomycin, 1× minimum essential medium non-essential amino acids, 1× vitamins, and 20 mM HEPES. All experiments were performed on cells between passages 6 and 15 to minimize culture-induced phenotypic drift.

*Cell Lines and General Culture Conditions*

HCT116 cells lacking all LRRC8 gene expression (8A through 8E all knocked out) were provided by Dr. Jerod Denton (Vanderbilt University). LRRC8A KO HEK293 cells were a gift from Dr. Rajan Sah, Washington University. HEK293 cells were grown in DMEM and HCT116 cells in McCoy's 5A medium, both supplemented with 10% FBS and 1% penicillin-streptomycin. All cell lines were maintained at 37°C in a humidified 5% $CO_2$ incubator. For signaling experiments, cells were serum-starved in medium for 2-3 hours prior to stimulation. The VRAC inhibitors carbenoxolone, DCPIB, and tamoxifen were used at final concentrations of 100 μM, 10 μM, and 10 μM, respectively, with a 30-minute preincubation.

*Redox Biology and Oxidative Stress Assays*

Superoxide Influx Measurement by DHE-HPLC: Superoxide influx in HEK293 cells was quantified by measuring the formation of the superoxide-specific product 2-hydroxyethidium (2-OH-$E^+$) using HPLC with fluorescence detection. Cells were seeded in 6-well plates and grown to ~90% confluence. After overnight serum starvation, cells were washed and loaded with 5 μM dihydroethidium (DHE) in phenol red-free HBSS for 30 minutes at 37°C. The DHE solution was removed and cells were washed three times before exposure to an extracellular superoxide-generating system consisting of 100 μM xanthine and 10 mU/mL xanthine oxidase (X/XO) and 500U/ml catalase for 15 minutes. The reaction was stopped by placing plates on ice and washing with ice-cold PBS. Cells were scraped in 200 μL of acidified acetonitrile, sonicated on ice and centrifuged at 16,000 x g for 10 minutes at 4°C. The supernatant was injected onto a C18 reverse-phase column (e.g., Agilent Zorbax Eclipse Plus) and analyzed on an HPLC system equipped with a fluorescence detector set to Ex/Em 480/580 nm. Peak areas corresponding to the 2-OH-$E^+$ standard were integrated and the values were normalized to total protein content of the well, as determined by a BCA protein assay (Thermo Fisher).

Live-Cell Confocal Imaging of superoxide: VSMCs were seeded on 35 mm glass-bottom dishes (MatTek). To measure mitochondrial superoxide, cells were loaded with ROSstar 550 (LI-COR Biosciences, 10 μM) indicator for 30 minutes at 37°C in HBSS. Cells were imaged live on a Zeiss LSM 880 laser-scanning confocal microscope equipped with an environmental chamber (37°C, 5% CO2). Images were acquired using a 561 nm laser for excitation, with emission collected between 570–620 nm. All acquisition parameters (laser power, gain, pinhole) were kept constant across all groups within an experiment.

For flow-cytometry assays, primary aortic VSMCs were grown to ~70–80% confluence, washed with PBS, and detached with trypsin. Cells were counted, and $2.5 \times 10^5$ cells per sample were aliquoted and resuspended in serum-free VSMC medium. Cells were then incubated with MitoSOX Red (Thermo Fisher, #M36008) (2.5 μM), TMRE (Thermo Fisher, #T669) (50 nM), or

ROSstar550 (10 μM) for 30 minutes at 37°C in the dark. After staining, cells were washed once with PBS.

For acquisition, TMRE-stained samples were resuspended in serum-free VSMC medium, whereas MitoSOX- and ROSstar550-stained samples were resuspended in FACS buffer (2% FBS in PBS). All steps after dye addition were performed protected from light. Samples were run immediately on a BD FACSMelody flow cytometer (BD Biosciences). Forward and side scatter were used to gate single, DAPI negative (viable) cells and exclude debris and doublets; dye-negative controls were used to define positive gates for each fluorophore.

For each sample, 10,000 events were collected. MitoSOX, TMRE and ROSstar 550 fluorescence were exciting using the 488 nm laser. Fluorescence intensity values for the relevant channels were exported and analyzed using the Floreada.io online platform. For each biological replicate, the median (or mean) fluorescence intensity of dye-positive cells was used for statistical analysis. These values are displayed in the figures as violin plots which represent the pooled distribution of single-cell dye intensity across 3 independent experiments.

*Lipid Peroxidation*

Malondialdehyde (MDA), a stable product of lipid peroxidation, was quantified using a thiobarbituric acid reactive substances (TBARS) assay kit (Apexbio, #K739-100). Cell lysates were prepared in PBS by sonication. Lysates reacted with thiobarbituric acid (TBA) at 95°C for 60 minutes. After cooling, the resulting pink TBARS adduct was measured calorimetrically by reading the absorbance at 532 nm on a plate reader. MDA concentration was calculated from a freshly prepared standard curve and normalized to the total protein content of the lysate.

*Glutathione (GSH/GSSG) Ratio*

The ratio of reduced (GSH) to oxidized (GSSG) glutathione, a key indicator of cellular redox state, was determined using the GSH/GSSG-Glo Luminescence Assay (Promega, #V6611) as per the manufacturer's protocol. For total glutathione (GSH + GSSG), cell lysates were incubated with a reagent containing a luciferin derivative and glutathione S-transferase (GST). For specific GSSG measurement, a parallel set of samples was pre-treated with N-ethylmaleimide to scavenge and block free GSH before lysis. Luminescence, which is proportional to the amount of GSH present, was measured on a plate reader. Concentrations were determined from a standard curve and the GSH/GSSG ratio was calculated and normalized to protein content.

*Lentiviral Luciferase Reporter Assays*

To quantitate transcription factor activity, primary VSMCs were transduced with custom-made lentiviral particles encoding firefly luciferase reporters under the control of response elements for NF-κB (pLV-κB-luc), Hif-1α (pLV-HRE-luc), or Nrf2 (pLV-ARE-luc). All reporters co-expressed Green Fluorescent Protein (GFP) to monitor infection efficiency.

Lentivirus Production: To produce lentiviral particles, HEK293T cells were co-transfected with the respective reporter plasmid, a packaging plasmid (psPAX2), and an envelope plasmid (pMD2.G) using Polyethylenimine (PEI) as the transfection reagent. Conditioned media containing the viral particles was harvested 48 hours post-transfection, clarified by centrifugation, and filtered through a 0.45 μm syringe filter.

Primary VSMCs were isolated from three WT and three LRRC8A KO mice, cultured, and then pooled in equal numbers by genotype before infection to ensure biological representation. The pooled VSMCs were transduced by incubation with the viral supernatant, supplemented with 8 μg/mL polybrene, for 48 hours. Following transduction, cells were serum-starved and then subjected to specific stimuli. For HIF-1α activation, cells were either treated with 300 μM cobalt chloride ($CoCl_2$) for 6 hours or placed in a modular hypoxia chamber flushed with a gas mixture of 1% $O_2$, 5% $CO_2$, and balanced $N_2$ for 24 hours. For NF-κB, cells were treated with 10 ng/mL TNFα for 6 hours. After stimulation, cells were lysed in Passive Lysis Buffer (Promega). Luciferase activity was measured using the Luciferase Assay System (Promega) on a luminometer. A multi-step normalization was performed to ensure accurate comparison between genotypes. First, raw luciferase activity (Relative Light Units) was normalized to the total protein concentration of the lysate, determined by a BCA assay. Because initial tests revealed that KO VSMCs had a higher transduction efficiency than WT cells, further correction was required. The relative level of transduction was quantified by performing a Western blot on the cell lysates to measure the amount of co-expressed GFP, which was then normalized to α-tubulin. The final reported luciferase activity was adjusted by this GFP/tubulin ratio to accurately reflect the transcriptional activity per successfully transduced cell, thereby correcting for variations in infection rate.

*Bulk RNA Sequencing and Bioinformatic Analysis*

Sample Preparation and RNA Isolation: Total RNA was extracted from three independent biological replicates of primary VSMCs derived from 3 WT and 3 LRRC8A KO mice. Cells grew to ~80% confluence before being lysed directly in the culture dish. RNA was isolated using the QIAGEN RNeasy Plus Mini Kit, which includes a genomic DNA eliminator column, followed by an additional on-column DNase I digestion step to ensure complete removal of genomic DNA. The quantity and purity of the extracted RNA were assessed using a Nanodrop spectrophotometer (A260/280 and A260/230 ratios > 1.9). The integrity of the RNA was rigorously evaluated using an Agilent 2100 Bioanalyzer; only samples with an RNA Integrity Number (RIN) of 9.0 or higher were carried forward for library preparation.

Library Preparation and High-Throughput Sequencing: Stranded, poly(A)-selected mRNA libraries were prepared from 1 μg of total RNA per sample using an Illumina-compatible library preparation kit with unique dual indices for multiplexing. The quality and size distribution of the final libraries were confirmed using the Agilent Bioanalyzer. The libraries were then pooled and sequenced on an Illumina NovaSeq 6000 platform to generate paired end reads (2 x 150 bp), achieving a sequencing depth of at least 30 million read pairs per sample. Bioinformatic analysis of the raw sequencing data was performed using Partek® Flow® software (Partek Inc., St. Louis, MO). The standardized pipeline within Partek included: 1) Quality assessment and trimming of raw FASTQ reads to remove adapters and low-quality bases; 2) Alignment of the clean reads to the mouse reference genome (GRCm39/mm39) using the STAR aligner; and 3) Quantification of reads to the gene level based on GENCODE annotation. The resulting gene count matrix was then used for differential gene expression analysis compared KO vs. WT VSMCs. This analysis was performed within Partek Flow using its implementation of the DESeq2 algorithm, which normalizes for library size and models variance. The output, including $\log_2$ fold change, p-values, and false discovery rates (FDR), was exported for further analysis. Genes with an FDR < 0.05 were considered significantly differentially expressed. The list of differentially expressed genes was used for several downstream analyses and visualizations. To interpret biological significance, gene lists were uploaded to the Enrichr web platform for functional enrichment analysis to identify enriched KEGG pathways, Gene Ontology (GO) terms and to perform protein-protein interaction (PPI) hub analysis. Gene-disease associations were explored using the DisGeNET database. To visualize the results of the differential expression analysis, volcano plots were generated using the web-based tool VolcaNoseR. Hierarchically clustered heatmaps illustrating the expression patterns of selected gene sets were created using both the Morpheus software from the Broad Institute and the Heatmapper web server. To compare gene lists and identify common or unique genes between different conditions, Venn diagrams were generated using the Venny 2.1 online tool.

The raw and processed sequencing data have been deposited in the NCBI Gene Expression Omnibus (GEO) public repository and are available under the accession number GSEXXXXXX.

*Gene-expression qPCR*

For gene-expression analyses, total RNA was isolated from primary aortic VSMCs (WT and LRRC8A-KO) using a column-based purification kit according to the manufacturer's instructions. RNA purity and concentration were assessed spectrophotometrically, and only samples with A260/280 ≈ 2.0 were used. One microgram of DNase-treated RNA was reverse-transcribed using a standard reverse transcriptase and a mixture of random hexamers and oligo(dT) primers to generate cDNA.

Quantitative PCR was performed in 96-well plates using a SYBR Green–based master mix on a real-time PCR system (e.g., CFX96, Bio-Rad). Each 10–20 µL reaction contained 1–2 µL cDNA and 0.2–0.4 µM of each primer. Primers for Rps4l, Hk2, Pfkp, and Nrf2 target genes (Gpx3, Gsta3, Hmox1) are listed in Supplemental Table 1. All samples were run in technical triplicate, and no-template controls were included on every plate. Melt-curve analysis was used to confirm amplification of a single specific product. Ct values were normalized to the housekeeping gene Gapdh, which was stable across experimental conditions. Relative mRNA abundance was calculated using the $2^{-\Delta\Delta Ct}$ method, using WT normoxia as the calibrator for hypoxia/$CoCl_2$ experiments and the mean WT control for other comparisons, as indicated in the figure legends.

*Metabolism*

Mitochondrial oxygen consumption and glycolysis assays

Mitochondrial respiration and glycolytic flux were quantified using an XF extracellular flux analyzer (Seahorse XFe96, Agilent). Primary aortic VSMCs from WT and LRRC8A-KO littermates were seeded in XF96 cell-culture plates at $1.2 \times 10^4$ cells/well and allowed to adhere overnight in complete VSMC medium. On the day of assay, cells were washed and incubated in bicarbonate-free Seahorse assay medium, supplemented as specified below, and equilibrated for 45–60 minutes in a non-$CO_2$ incubator.

For the glycolysis stress test, cells were maintained initially in glucose-free assay medium, and the extracellular acidification rate (ECAR) was recorded at baseline, after addition of glucose (10 mM), after oligomycin (1 mM), and following 2-deoxy-D-glucose (2-DG, 50 mM). These sequential injections allowed calculation of basal glycolysis, glycolytic capacity, and glycolytic reserve according to the manufacturer's instructions.

For the mitochondrial stress test, assay medium was supplemented with 10 mM glucose, and oxygen consumption rate (OCR) was measured at baseline, after oligomycin (1 mM), carbonyl cyanide-p-trifluoromethoxy phenylhydrazone (FCCP, 1 mM), and antimycin A/rotenone (0.5 mM each). From these traces we derived basal respiration, ATP-linked respiration, maximal respiratory capacity, non-mitochondrial respiration, and proton leak.

After each run, plates were inspected to confirm even cell distribution and absence of edge artifacts. ECAR and OCR values were normalized to cell number or protein content per well, as indicated in the figure legends. For each genotype, data represents the mean of independent biological preparations (multiple wells per run), and statistical comparisons were performed as described in the main Methods.

The ADP/ATP ratio, an indicator of cellular energy status, was measured using the ADP-Glo Kinase Assay (Promega, #V9101). The $NADP^+$/NADPH ratio, a key indicator of cellular reductive capacity, was measured using the NADP/NADPH-Glo Assay (Promega, #V6611). For both assays, VSMCs were seeded in 96-well plates and assays were performed according to the manufacturer's protocols. In brief, cells were lysed to stop enzymatic activity and stabilize nucleotides. Specific detection reagents were then added to generate a luminescent signal proportional to the amount of ADP or $NADP^+$/NADPH. Luminescence was measured on a plate reader. Ratios were calculated after determining absolute concentrations from standard curves and were normalized to protein content per well.

*Cellular viability Assays*

VSMC viability was assessed using the colorimetric MTT (3-(4,5-dimethylthiazol-2-yl)-2,5-diphenyltetrazolium bromide) assay. Cells were seeded at a density of 5,000 cells/well in 96-well plates. At indicated time points (0h, 24h, 48h, 72h), MTT reagent was added to each well to a final concentration of 0.5 mg/mL and incubated for 4 hours at 37°C to allow for the formation of formazan crystals. The medium was then removed, and the formazan crystals were solubilized in 100 µL of DMSO. Absorbance was measured at 570 nm with a reference wavelength of 650 nm using a microplate reader.

Cell Migration (Scratch Wound Healing Assay): VSMC migration was quantified using the IncuCyte S3 Live-Cell Analysis System (Sartorius). Cells were seeded in 96-well ImageLock plates and grown to a confluent monolayer. A uniform 700-800 μm scratch was made simultaneously in all wells using the 96-pin IncuCyte Wound Maker tool. Wells were washed twice with PBS to remove detached cells and debris and then refilled with migration medium (DMEM with 0.5% FBS) with or without 10 ng/mL TNFα. The plate was placed in the IncuCyte system and phase-contrast images of the wounds were acquired every 3 hours for 72 hours. The IncuCyte S3 software automatically measured the average wound width (in μm) at each time point. Migration rate was determined from the slope of the linear phase of wound closure.

Senescence-Associated β-Galactosidase (SA-β-Gal) Activity: VSMCs were seeded at a density of 1×105 cells per well in 6-well plates and cultured for 6 days in medium containing either 10% FBS or 0% FBS (serum-free) to induce stress. SA-β-Gal activity was then measured using the quantitative Cellular Senescence Activity Assay Kit (Thermo Fisher, #CBA-230), which utilizes the fluorogenic substrate 4-methylumbelliferyl β-D-galactopyranoside (4-MUG). Cell lysates were incubated with the substrate and the fluorescence of the hydrolyzed product was measured on a plate reader (Ex/Em 365/460 nm). Activity was normalized to total protein content.

*Relative Telomere Length Measurement by qPCR*

Genomic DNA (gDNA) was isolated from passage-matched WT and LRRC8A-KO VSMCs using the DNeasy Blood & Tissue Kit (QIAGEN). The concentration and purity of the gDNA were determined, and 5-10 ng of DNA was used per reaction. Relative telomere length was measured using the quantitative PCR method described by Cawthon, which calculates the ratio of telomere repeats (T) to a single-copy gene (S). Two separate qPCR reactions were run for each sample in triplicate on a Bio-Rad CFX96 Real-Time PCR Detection System using PowerUp SYBR Green Master Mix (Applied Biosystems): one amplifying the telomere repeats and the other amplifying the single-copy reference gene Rplp0 (also known as 36B4). Raw $C_t$ values from technical replicates with a standard deviation > 0.25 were excluded from the analysis. The relative telomere length (T/S ratio) was calculated using the $2^{\Delta\Delta}C_t$ method, where $^{\Delta}C_t$ was first determined as ($C_t^{Telomere} - C_t^{36B4}$). To normalize results across different runs, this $^{\Delta}C_t$ was then compared to a calibrator sample (a master pool of WT gDNA) included on every plate to derive the $^{\Delta\Delta}C_t$.

*Western Blotting*

Proteins were extracted from cultured primary VSMCs, whole aortae (thoracic and abdominal aortae) or mesenteric arteries. Aortae and mesenteric arteries were first cleaned of adherent fat and connective tissue in ice-cold physiological salt solution (PSS), flash-frozen in liquid nitrogen, and stored at -80°C. Prior to lysis, frozen tissues were pulverized to a fine powder using a liquid nitrogen-chilled mortar and pestle. The tissue powder or cell pellets were then lysed on ice for 1 hour in a phosphoprotecting lysis buffer (PPLB) containing 50 mM Tris-base (pH 7.4), 150 mM NaCl, 1 mM EDTA, 1 mM DTT, 10% glycerol, 1% Triton X-100, 0.1% Na-deoxycholate, 0.1% SDS, and a cocktail of phosphatase and protease inhibitors (10 mM β-glycerophosphate, 20 mM p-nitrophenyl phosphate, 2 mM sodium pyrophosphate, 1 mM $Na_3VO_4$, 5 mM NaF, 10 μg/mL aprotinin, and 1 mM PMSF). The resulting lysates were clarified by centrifugation at 20,000 x g for 20 minutes at 4°C. The protein concentration of the clarified supernatant was determined using a BCA protein assay. For analysis, 20-50 μg of total protein per sample was denatured, resolved by SDS-PAGE, and transferred to nitrocellulose membranes. The membranes were blocked for 1 hour at room temperature with Odyssey Blocking Buffer (LI-COR Biosciences, Lincoln, NE) and then incubated overnight at 4°C with primary antibodies diluted in Tris-buffered saline with 0.1% Tween-20 (TBS-T). A comprehensive list of all primary antibodies and their sources is provided in the supplementary materials. Following primary antibody incubation, membranes were washed in TBS-T and incubated with corresponding fluorescently conjugated secondary antibodies for 2 hours at room temperature. The fluorescent signals were detected and quantified using an Odyssey Infrared Imaging System and its accompanying Image Studio software (LI-COR Biosciences). Band intensities for target proteins were normalized to a loading control (e.g., α-tubulin or GAPDH) to correct for variations in protein loading. For the lentiviral reporter assays, this method was also used to quantify GFP expression, which was normalized to α-tubulin to correct for differences in transduction efficiency between samples.

*Development of atherosclerotic animals*

$ApoE^{-/-}$ $LRRC8A^{-/-}$ (double KO) mice were created by breeding $ApoE^{-/-}$ (Jackson Laboratory) and VSMC-specific LRRC8A KO. In atherosclerosis study, all mice were $ApoE^{-/-}$ background. LRRC8A WT, $LRRC8A^{+/-}$ and LRRC8A KO mice were fed with high fat diet (42% Fat, TD88137, Harlan Teklad) for 15 weeks after weaning to induce atherosclerosis. All procedures were performed in accordance with the *Guiding Principles in the Care and Use of Animals*, approved by the Vanderbilt University Institutional Animal Care and Use Committee. The animals were housed on a 12-hour light/dark cycle and fed with water *ad libitum*. For the study, mice were euthanized by placement in a sealed chamber with 100% $CO_2$ provided at a flow rate of 2 L/min for 3 min. At that time, mice were monitored till they had > 1 min of respiratory cessation and blood was collected from heart. PCR analysis of tail-tip DNA was used to identify LRRC8A WT, LRRC8A$^{+/-}$, or LRRC8A KO with ApoE$^{-/-}$ mice through TransnetYX, Inc.

*Assessment of atherosclerosis and lipid measurement*

Atherosclerotic lesion area was examined with Oil Red-O in whole aorta and quantified by *en face* analysis. Whole aorta was dissected from heart to iliac artery branch point and cleaned in cold physiological salt solution (PSS; 130 mmol/L NaCl, 4.7 mmol/L KCl, 1.18 mmol/L $KH_2PO_4$, 1.18 mmol/L $MgSO_4 \cdot 7H_2O$, 1.56 mmol/L $CaCl_2 \cdot 2H_2O$, 14.9 mmol/L $NaHCO_3$, 5.6 mmol/L glucose, and 0.03 mmol/L EDTA). The cleaned and opened aorta was fixed in 4% paraformaldehyde in PBS overnight in 4 °C. Fixed aorta was washed with water and stained with Oil Red-O solution (#26503-02, Electron Microscopy Sciences, Hatfield, PA) for 2 hours. The aorta was subsequently immersed in 60% isopropyl alcohol for 2 minutes and washed with water 3 times. Atherosclerotic lesion area was stained with red and quantitated in ImageJ software.

Collected blood samples were left in room temperature for 2 hours and serum samples were isolated. Samples were sent to Vanderbilt Lipid Core (Vanderbilt University) to measure total cholesterol and triglyceride. Total cholesterol above 2000 mg/dL was indicated as 2000 mg/dL.

*Cholesterol uptake assay*

Red-orange fluorescent Dil dye labeled oxLDL (Dil-oxLDL; #L34358, Thermo Fisher Scientific) or non-oxidized LDL (Dil-LDL; #L3482, Thermo Fisher Scientific) was used to determine cholesterol uptake. VSMCs were treated with Dil-oxLDL or Dil-LDL (10 µg/mL) for

4 hours, washed with PBS, and then fixed in 4 % paraformaldehyde in PBS. The image of oxLDL or LDL uptake was captured by confocal microscopy and z-projection images were analyzed with ImageJ software.

*Evaluation of vascular function*

First- or second-order branches (~100 µM inner diameter) of the superior mesenteric arteries were excised, cleaned of fat and connective tissue and cut into 1 to 2 mm length-rings in an ice-cold PSS. Mesenteric rings were mounted in wire myographs (Danish Myo Technology A/S, Aarhus, Denmark) containing warmed (37 °C) and oxygenated (95% $O_2$/5% $CO_2$) PSS and allowed to equilibrate for 45 min under a passive force of 2 - 2.5 mN. Arterial viability was assessed by stimulation with 120 mM KCl. After washing, rings were contracted with phenylephrine (PE, $5\times10^{-7}$ mol/L), followed by exposure to acetylcholine (ACh, $10^{-6}$ mol/L). A more than 70% relaxation response to ACh was taken as evidence of an intact endothelial layer. Endothelium-dependent relaxation was assessed on PE-contracted ($10^{-6}$ mol/L) rings by cumulative addition of ACh ($10^{-9}$ to $10^{-5}$ mol/L), while endothelium-independent relaxation was tested using sodium nitroprusside (SNP, $3\times10^{-9}$ to $10^{-6}$ mol/L). Contractile responses were assessed by cumulative exposure to PE ($10^{-9}$ to $3\times10^{-5}$ mol/L) or NE (norepinephrine; $10^{-9}$ to $3\times10^{-5}$ mol/L). Contractions were recorded as changes in tension (mN) from baseline, expressed as a percentage of the response to 120mM KCl. Relaxation was expressed as a percentage of the stable contraction produced by PE in each ring immediately prior to the first dose of vasodilator. Return to the tension recorded before addition of PE was considered as 100% relaxation.

*Statistical Analysis*

All quantitative data are expressed as the mean ± standard error of the mean (SEM). The variable 'n' represents the number of individual animals or the number of independent biological replicates for experiments performed in cultured cells. All statistical analyses were performed using GraphPad Prism (Version 9.0, GraphPad Software, San Diego, CA).

The specific statistical test used was chosen based on the experimental design. For direct comparisons between two groups (e.g., WT vs. KO), an unpaired, two-tailed Student's T-test was employed. For data that were normalized to a control group set to a value of 1.0, a one-sample t-test was used to determine if the experimental group was significantly different from the control. For experiments involving three or more groups, statistical differences were assessed by a one-way or two-way Analysis of Variance (ANOVA). If the ANOVA result was significant, a Tukey's multiple comparisons test was performed for post hoc analysis to identify specific differences between group means.

For vascular reactivity studies, concentration-response curves were generated using nonlinear interactive regression analysis to determine two key pharmacological parameters: the maximal effect ($E_{max}$) and the $EC_{50}$ (the molar concentration of an agonist required to produce 50% of the maximum response).

In all cases, the probability value of $p < 0.05$ was statistically significant.
